# Inhibition of multidrug-resistant *Staphylococcus aureus* by commensal bacterial species from the human nose

**DOI:** 10.64898/2026.02.27.708567

**Authors:** Anaëlle Fait, Daniel C. Angst, Valeria Stylianou, Laura Brülisauer, Silvio D. Brugger, Alex R. Hall

**Affiliations:** Department of Environmental Systems Science, Institute of Integrative Biology, ETH Zürich, Zürich, Switzerland; Department of Infectious Diseases and Hospital Epidemiology, University Hospital Zürich, University of Zürich, Zürich, Switzerland

## Abstract

*Staphylococcus aureus* is an important human pathogen. With the growing threat of antimicrobial resistance, exemplified by methicillin-resistant *S. aureus* (MRSA) infections requiring last-line antibiotics, alternative strategies to combat *S. aureus* and MRSA colonisation are needed. One promising direction is to harness interactions between *S. aureus* and commensal bacteria resident in the nasal passage. However, it remains unclear which bacterial species or combinations are most effective at inhibiting *S. aureus*, and whether this inhibition is affected by resistance to last-line antibiotics. Here, we address this gap using systematic *in vitro* experiments to identify commensal species or combinations that inhibit MRSA, including strains resistant to key last-resort therapies. In a combinatorial screen, we detected widespread inhibition of MRSA by commensal species, with inhibitory strength dependent on species identity and community composition. Staphylococcal commensal species produced the strongest MRSA inhibition, but we achieved similarly strong inhibition by combining *Corynebacterium* species with *Dolosigranulum pigrum*. This was linked to positive growth interactions between commensals. Importantly, we observed similar levels of MRSA inhibition by commensal species when we tested MRSA strains resistant to key anti-MRSA antibiotics, including clinical isolates. Inhibition was sustained during serial passaging experiments, with minimal MRSA adaptation upon prolonged exposure. Our results show defined combinations of nasal commensals can robustly inhibit MRSA, including strains resistant to last-resort antibiotics. This supports the potential of microbiota-based therapies to prevent or eliminate *S. aureus* colonisation from the human nose, including in the context of antibiotic resistance.

## Introduction

An estimated 1.14 million people died from antibiotic-resistant bacterial infections in 2021, with methicillin-resistant *Staphylococcus aureus* MRSA alone accounting for 130 000 deaths (Naghavi et al., 2024). MRSA isolates are resistant to most β-lactam antibiotics (Hartman & Tomasz, 1984), limiting treatment options and necessitating alternative strategies. One promising approach is through manipulation of the nasal microbiome (Ghosh et al., 2019; Panwar et al., 2021). The nose is the main ecological niche for *S. aureus* and nasal carriage is a risk factor for infections, which are most often initiated by the colonising strain (Clarridge et al., 2013; von Eiff et al., 2001; Wertheim et al., 2005). The nasal cavity hosts complex bacterial communities that can naturally inhibit pathogens, including *S. aureus* (Adolf & Heilbronner, 2022; Brülisauer et al., 2026; Escapa et al., 2018; Krismer et al., 2014). However, to translate into effective interventions, it is critical to identify which species or combinations of commensals drive competitive interactions against *S. aureus*. Pairwise *in vitro* experiments and epidemiological studies have shown some commensals, such as *Dolosigranulum pigrum*, *Corynebacterium* spp., and *Staphylococcus lugdunensis*, can inhibit *S. aureus* or negatively correlate with *S. aureus* carriage (Brugger et al., 2020; Hardy et al., 2020; Liu et al., 2015a; Wos-Oxley et al., 2010; Zipperer et al., 2016). However, these approaches provide an incomplete picture. Pairwise experiments overlook potential interactions between different commensal species, and epidemiological data are often confounded by host and environmental variables. Thus, it remains incompletely understood which species, individually or in combinations, provide the most robust inhibition of MRSA. Addressing this challenge would advance our ability to design targeted microbiota-based interventions.

A second key challenge in developing microbiota-based approaches to inhibit MRSA is to account for emerging resistance against other antibiotics. MRSA infections are typically treated with last-resort antibiotics such as daptomycin, vancomycin, and teicoplanin, but the emergence of resistance to these drugs is further limiting therapeutic options (Parsons et al., 2025). Resistance to these antibiotics may alter susceptibility of MRSA to inhibition by other microbes if there are shared molecular targets. For example, both vancomycin and nisin (a bacteriocin) target lipid-II inside the cell wall of *S. aureus*. Supporting the potential for cross-resistance, vancomycin resistance has been demonstrated to arise and be maintained upon exposure to bacterial competition and/or exposure to bacteriocins (Fait et al., 2023; Koch et al., 2014). More generally, antibiotic adaptation in *S. aureus* and in other species can lead to pleiotropic changes in other phenotypes, including susceptibility to other antibiotics (Pál et al., 2015), population growth (Qi et al., 2014), stress responses (Paulander et al., 2009), and production of compounds inhibitory to other species (Quinn et al., 2022). In *E. coli*, pleiotropic changes in metabolism associated with antibiotic resistance were linked to increased competitive ability relative to other microbes and colonisation success in the mammalian intestinal tract (Connor et al., 2023). Antibiotic adaptation in *S. aureus* may therefore lead to unpredictable changes in susceptibility to inhibition by other microbes, but this relationship is not yet well characterized. Addressing this gap in our knowledge is important, because the value of microbiota-based strategies to combat MRSA will be particularly high in situations where conventional antibiotic therapy is compromised by resistance.

We studied the interaction between MRSA isolates and bacterial species representative of the human nose microbiome (*Cutibacterium acnes*, *Corynebacterium accolens, Staphylococcus lugdunensis, Corynebacterium tuberculostearicum, Dolosigranulum pigrum, S. aureus* and Neisseria sicca; **Table S1**) (Escapa et al., 2018). To get closer to the spatial environment in the nose and enable direct observation of commensal-mediated effect on MRSA growth, we developed an experimental set-up on semi-solid surface for interaction screening. This system was designed to recapitulate key features of early-stage nasal colonisation under controlled conditions. To identify species and combinations that inhibit MRSA colonisation, we assembled all possible combinations of species and assessed colonisation resistance to MRSA strain JE2, a clinically relevant and widely studied community-associated USA300 isolate (Fey et al., 2013). We then tested commensal inhibition of MRSA strains with additional resistance to key anti-MRSA antibiotics, including clinical isolates. Our results show inhibition of MRSA strain JE2 by all tested commensal species, but to varying degrees. Intra-specific competition between commensal *S. aureus* (isolated from healthy nose) and JE2 yielded strong inhibition, yet a similar level of inhibition was achieved by combining *Corynebacterium* species with *D. pigrum*. Furthermore, inhibition was similarly effective against MRSA strains resistant to key anti-MRSA antibiotics, including clinical isolates. These findings highlight the potential of leveraging specific nasal commensal bacteria as a microbiota-based strategy to combat MRSA colonisation and infection, even in the context of resistance to last-resort antibiotics.

## Results

### Commensal species inhibit MRSA in colonisation assays

To determine which commensal species or combinations were effective at inhibiting MRSA, we screened all 128 possible combinations of the seven commensal species for their ability to limit growth of MRSA strain JE2. We developed an experimental set-up for colonisation on semi-solid media, first cultivating lawns of resident commensal bacteria, before inoculating JE2 at a relatively low density (see Methods). We did this at 34°C, the average temperature in the nasopharynx (Keck et al., 2000). We observed inhibition of JE2 by commensal species in several treatments (**Figure 1A**). Treatments containing Staphylococci (either or both of *S. aureus* and *S. lugdunensis*) provided the strongest inhibition. This effect was similarly strong across different combinations containing Staphylococci and Staphylococci alone (**Figure 1A and S1**), reducing final JE2 abundance from, on average, 9.96 ± 0.40 log_10_CFU/mL in the control treatment to 5.48 ± 0.46 log_10_CFU/mL in treatments with Staphylococci. However, excluding the treatments containing Staphylococci, we observed significant inhibition of JE2 in several other combinations comprising other commensal species (**Figure S1**).

**Figure 1.**
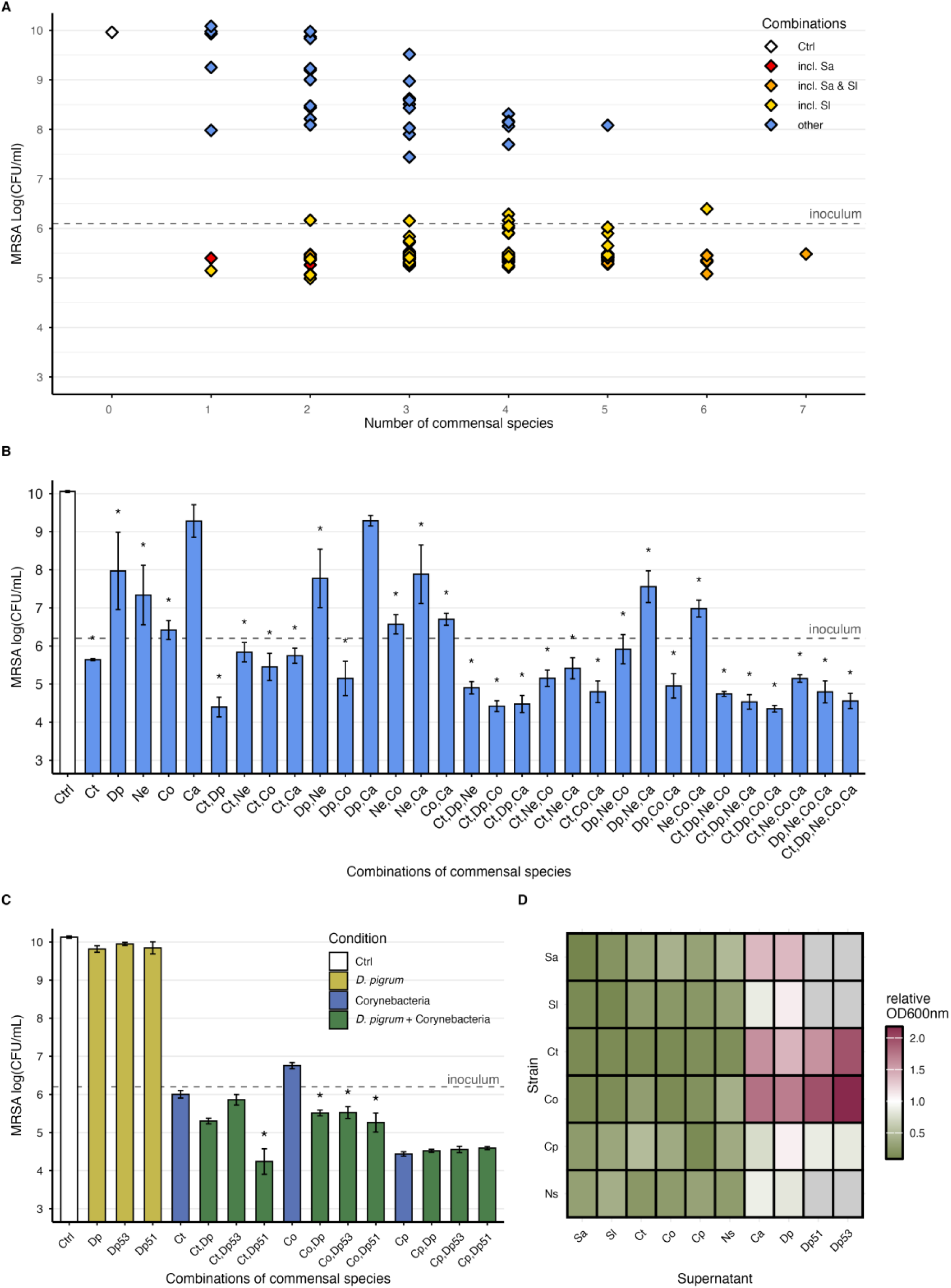
Inhibition of MRSA strain JE2 in colonisation assays. **A.** Growth of JE2 measured as log(CFU/mL) when co-cultured with combinations of seven commensal species on BHIA. Combinations that include *S. aureus*, *S. lugdunensis*, or both, are represented in red, yellow, and orange, respectively. Points represent the mean of four biological replicates. Growth of JE2 in each individual commensal combination is shown in Figure S1. **B.** Growth of JE2 measured as log10(CFU/mL) when co-cultured with combinations of commensal species on BHIA + 1% Tween 80. Bars represent mean ± SE for four biological replicates. Statistical comparisons were performed using Dunnett’s test. Significance: \**padj* ≤ 0.05. **C.** Growth of JE2 measured as log10(CFU/mL) when co-cultured with combinations of *Corynebacterium* species and *D. pigrum* isolates on BHIA + 1% Tween 80. Bars represent mean ± SE. Pairwise comparisons between each combination treatment (Corynebacteria + *D. pigrum*) and the corresponding *Corynebacterium* single treatment (Ct, Co, Cp) were performed using Dunnett’s test. Significance: \**p* ≤ 0.05. **D.** Growth of commensal strains in supernatants derived from culture of other species. Growth is expressed as relative OD_600nm_ at 24 h, calculated by dividing growth in strain-derived supernatant by growth in control conditions (uninoculated wells). Grey cells indicate missing data. Data for *D. pigrum* as receiver are not included due to its lack of growth in pure cultures in liquid (see Table S3). Ctrl: no commensal, Sa: *S. aureus*, Se: *S. lugdunensis*, Ct: *C. tuberculostearicum*, Dp: *D. pigrum* (strains Dp, Dp53 and Dp51), Ns: *N. sicca*. Co: *C. accolens*, Cp: *C. propinquum*, Ca: *Cutibacterium acnes*.

We suspected inhibition of JE2 in some treatment groups would be enhanced by improving the growth conditions for non-staphylococcal commensal species. In particular, *C. tuberculostearicum* and *C. accolens* are lipophilic species that depend on external lipids for growth as they lack a complete fatty acid biosynthesis pathway (Funke et al., 1997). This requirement likely drives their adaptation to lipid-rich environments, including sebaceous skin sites and the nasal cavity, where host-derived lipids are readily available. Tween 80 represents a source of fatty acids beneficial for growth of lipophilic species and other fastidious isolates (Popowitch et al., 2024) and we found supplementation with 1% Tween 80 improved growth of four out of six tested non-staphylococcal commensal species (**Figure S2**). When we tested for JE2 inhibition by commensals and combinations that did not include Staphylococci under these improved conditions, we observed inhibition in the majority of treatment groups (**Figure 1B**). Inhibition was on average strongest in treatment groups that included *C. tuberculostearicum* and/or *C. accolens*. In support of this interpretation, the presence/absence of these species accounted for 41% and 16% of the variance between treatments, respectively (ANOVA F_1,111_ = 268.018 and F_1,111_ = 105.385, p < 0.001; **Table S2**). *D. pigrum*, despite not significantly inhibiting JE2 as a single-species treatment, was a component of several inhibitory combinations (**Figure 1B**), and explained 7% of the variance among treatment groups (ANOVA, F_1,111_ = 46.676, p < 0.001; **Table S2**). In particular, treatment combinations including *D. pigrum* and either/both of *C. tuberculostearicum* and *C. accolens* provided strong JE2 inhibition. For example, the combination of *D. pigrum* and *C. tuberculostearicum* was sufficient to account for the strongest inhibition observed in this assay, with no further reduction in JE2 abundance upon addition of other species (**Figure 1B**). In these conditions, several treatments decreased the density of JE2 compared to the initial inoculum, suggesting killing rather than growth inhibition of JE2 (**Figure 1B**).

We next asked whether the strong inhibition observed with combinations of *D. pigrum* and either/both of *C. tuberculostearicum* and *C. accolens* applied to other species or strains within these families. To do this we tested combinations of three species of *Corynebacterium* with three strains of *D. pigrum*. For two of the three tested *Corynebacterium* species (*C. tuberculostearicum* and *C. accolens*), inhibition of JE2 was stronger in co-cultures with *D. pigrum* compared to the equivalent *Corynebacterium*-alone treatment (*padj* ≤ 0.05 for at least one of the three tested *D. pigrum* strains, by Dunnett’s test; **Figure 1C**). By contrast, while *C. propinquum* alone strongly inhibited JE2, co-culturing with *D. pigrum* did not amplify this effect (**Figure 1C**). As before, *D. pigrum* had little effect as a single-species treatment. In summary, commensal species from the human nasal microbiome were highly effective in suppressing growth of MRSA strain JE2. Strong inhibition was observed both in conditions including Staphylococci and in combinations of *D. pigrum* with *Corynebacterium* species.

### D. pigrum improves growth of Corynebacterium species

One possible explanation for the increased MRSA inhibition by combination treatments of *C. tuberculostearicum* and *D. pigrum* relative to the equivalent single-species treatments is mutual growth promotion. This could occur, for example, if one or both species modify the local abiotic conditions in a way that increases growth of the other. To test this, we cultured commensal species in supernatants derived from cultures of the other species, or control supernatant from uninoculated wells (see Methods). Six of the ten types of culture supernatant repeatably supported less growth than control supernatant (**Figure 1D, Table S3**), consistent with nutrient depletion or other mechanisms by which growth of one strain impairs growth of another. However, supernatant from *D. pigrum* strains improved growth of *Corynebacterium* species *C. tuberculostearicum* and *C. accolens* (**Figure 1D**). This specificity among supernatant types and receiving strains (supernatant × strain interaction in two-way ANOVA on log₂-transformed relative OD_600nm_: F_53,140_ = 5.04, P < 0.001), in particular among *Corynebacterium-D. pigrum* pairs, matched our prior observation that *D. pigrum*-*C. tuberculostearicum* and *D. pigrum*-*C. accolens* combinations, but not *D. pigrum-C. propinquum*, resulted in enhanced JE2 inhibition compared to respective *Corynebacterium* strains alone (**Figure 1C**). These findings suggest *D. pigrum* modified local abiotic conditions in a way that selectively promoted growth of certain *Corynebacterium* species, increasing MRSA inhibition (**Figure 1B**).

*Commensal treatments inhibit laboratory-adapted MRSA isolates resistant to anti-MRSA antibiotics* Genetic changes conferring antibiotic resistance can have wide-ranging pleiotropic effects on other phenotypes (Connor et al., 2023). Potentially, this could modify inhibition by other species. Therefore, we next tested whether commensal-species treatments that inhibited MRSA growth in our colonisation assays would also be effective against MRSA isolates with decreased susceptibility to key anti-MRSA antibiotics; vancomycin, teicoplanin and daptomycin. We generated three resistant isolates for each antibiotic by exposing JE2 to increasing concentrations of daptomycin and teicoplanin, then sequenced the resulting isolates (**Table S4 and S5**). Daptomycin is a lipopeptide antibiotic targeting the cell membrane. Accordingly, daptomycin-adapted isolates (DPC1, DPC2, DPC3) had mutations in *mprF* and *pgsA*, encoding proteins involved in cell membrane synthesis, commonly reported in daptomycin *S. aureus* adaptation (Freeman et al., 2024). Daptomycin adaptation is often paired with decreased susceptibility to glycopeptide antibiotics vancomycin and teicoplanin, targeting peptidoglycan synthesis, which was also the case for our resistant mutants (**Table S4**) (Reynolds, 1989). Our mutant isolate DPC2 also had a mutation in *vraT* targeting the three-component system VraTSR, involved in cell wall stress response and often mutated during glycopeptide adaptation (**Table S5**) (Boyle-Vavra et al., 2013). Teicoplanin-resistant isolates (TEI1, TEI2, TEI3) had mutations in genes involved in cell wall synthesis and cell wall stress response, namely in *walk* targeting the essential two-component system WalKR, and *vraT* (**Table S5**) (Dubrac et al., 2008; Boyle-Vavra et al., 2013). Mutant isolate TEI3 acquired a mutation in *rpoB*, encoding the RNA polymerase beta subunit. Mutations in *rpoB* in glycopeptide-adapted strains are commonly associated with increased cell wall thickness (Cui et al., 2010). We also used three JE2-derived isolates with decreased susceptibility to vancomycin from a previous study, with mutations in common vancomycin-adaptive genes and either collateral resistance or susceptibility to beta-lactam antibiotics (Fait et al., 2022; VAN1, VAN4 and VAN9). Most resistant isolates had slower growth compared to wild-type JE2 (**Figure S3**). In summary, the resistant isolates we used in this section carried representative and clinically relevant resistance mutations (Thitiananpakorn et al., 2020), including in genes shown elsewhere to confer pleiotropic effects on other phenotypes (Mishra et al., 2011).

We carried out colonisation assays with wild-type JE2 and these resistant isolates in the presence/absence of six different commensal-species treatments. As in our earlier assays, we observed strong inhibition with most commensal-species treatments relative to the control treatment, especially treatments including both *C. tuberculostearicum* and *D. pigrum*, but not *D. pigrum* alone (**Figure 2A**). Exposure to different commensal treatments had similar effects on growth of JE2-derived antibiotic-resistant isolates compared with wild-type JE2 for 8 of 9 resistant isolates (Condition × Resistance interaction in two-way ANOVA, tested separately for each resistant isolate vs. the wild-type - *p* > 0.05 for all isolates except DCP3, for which F_6,42_ = 4.96, Holm-Bonferroni-adjusted P < 0.001). DPC3 was more strongly inhibited than the wild-type in combinations of *C. tuberculostearicum* and *D. pigrum* as well as all species combined (Holm-adjusted post-hoc mutant vs WT contrasts within condition: Ct,Dp, *padj* < 0.001; Sa,Sl,Ct,Dp, *padj* < 0.001; **Figure 2A**). Together, these data indicate that resistance to anti-MRSA antibiotics did not reduce the effectiveness of commensals at inhibiting MRSA.

**Figure 2.**
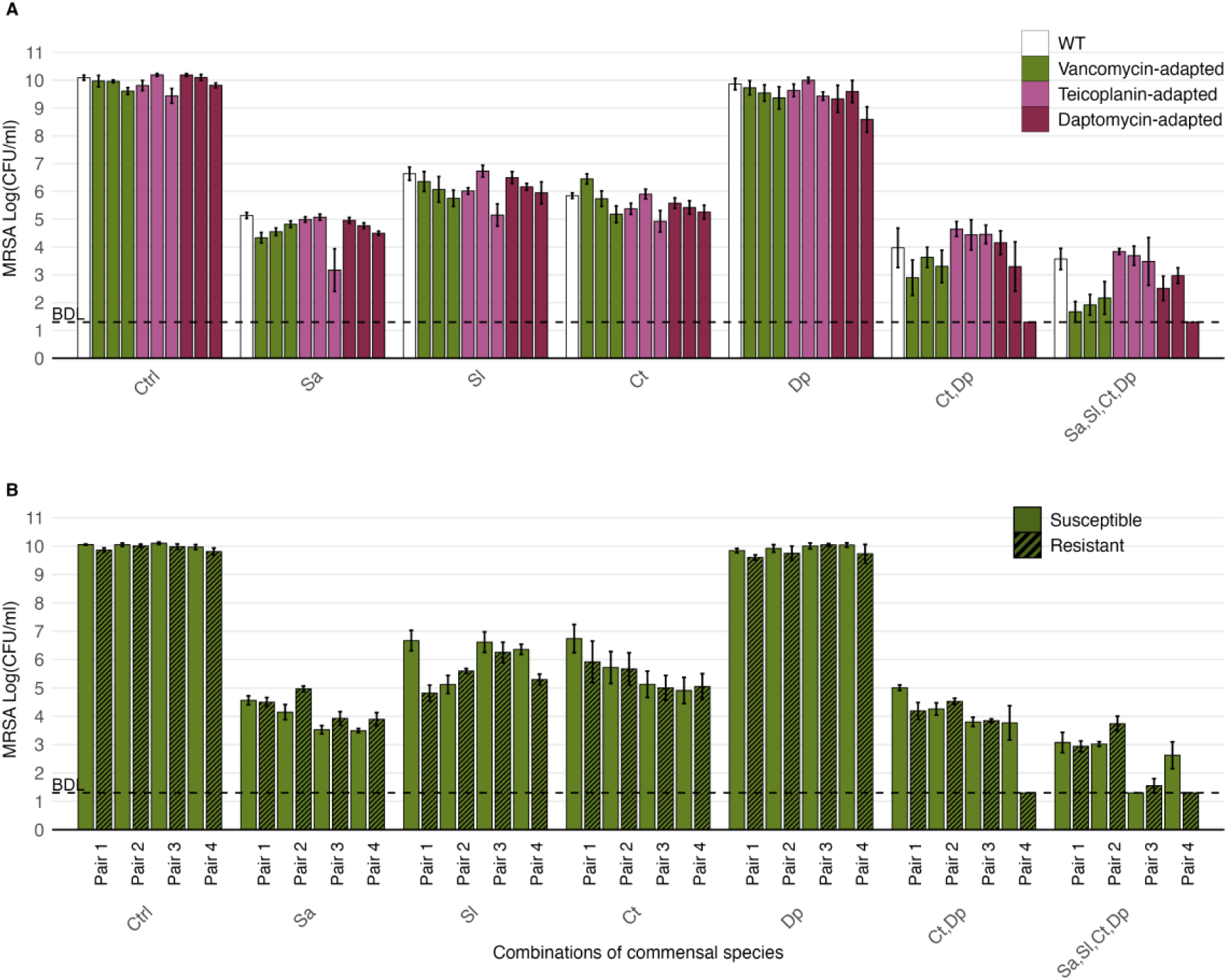
Commensal communities provide robust inhibition of MRSA isolates despite resistance to anti-MRSA antibiotics. **A.** Growth of JE2 and JE2-derived isolates with decreased susceptibility to vancomycin (VAN1, VAN4, VAN9), teicoplanin (TEI1, TEI2, TEI3) and daptomycin (DPC1, DPC2, DPC3) co-cultured with combinations of commensals, measured as log10(CFU/mL). Bars represent mean ± SE for four biological replicates. **B.** Growth of four pairs of MRSA clinical isolates susceptible or resistant to vancomycin co-cultured with combinations of commensals, measured as log10(CFU/mL). Bars represent mean ± SE of four biological replicates. BDL: below detection limit. Sa: *S. aureus*, Sl: *S. lugdunensis*, Ct: *C. tuberculostearicum*, Dp: *D. pigrum*.

### Commensal treatments inhibit clinical MRSA isolates resistant to anti-MRSA antibiotics

Clinical isolates are exposed to combined antibiotics and host selective pressures, which may modify their interaction with bacterial species from the host microbiome. We therefore tested four pairs of clinical isolates in our colonisation assay. Each pair comprised MRSA isolates retrieved before and after vancomycin treatment: JKD6000/JKD6001 (Pair 1), JKD6004/JKD6005 (Pair 2), JKD6009/JKD6008 (Pair 3) and JKD6052/JKD6051 (Pair 4) (Howden et al., 2006). JKD6001, JKD6005, JKD6008 and JKD6051 have been shown previously to have reduced susceptibility to vancomycin and teicoplanin, with JKD6051 also having rifampicin resistance (Fait et al., 2024; Howden et al., 2011), and to have key mutations in genes encoding the WalKR, VraTSR systems and *rpoB* (Howden et al., 2011). The pairs belong to the same clonal complex CC8 and sequence type ST239 (Howden et al., 2011). Across these clinical isolate pairs, we observed strong inhibition in multiple commensal treatments, particularly those including *C. tuberculostearicum* and *D. pigrum*, but not *D. pigrum* alone (**Figure 2B**). Inhibition was strongest for Pair 3 and Pair 4 when all four commensals were combined, resulting in complete eradication (below the detection limit) for two isolates. Resistant and susceptible isolates responded similarly to different commensal treatments in three of the four strain pairs (Condition × Resistance interaction in two-way ANOVA, tested separately for each strain pair - *p* > 0.05 for pairs 1-3; for pair 4, F_6,42_ = 5.51, p < 0.001). Specifically, the resistant isolate in pair 4 was even more strongly inhibited than the equivalent susceptible strain in treatments combining *C. tuberculostearicum* and *D. pigrum* (Holm-adjusted post-hoc susceptible vs resistant contrasts within condition: Ct,Dp, *padj* < 0.001; Sa,Sl,Ct,Dp, *padj* < 0.001; **Figure 2B**). Consistent with observations for DPC3 (**Figure 2A**), this isolate survived exposure to *D. pigrum* and *C. tuberculostearicum* individually but was eradicated when both species were combined. Overall, isolates with reduced susceptibility to vancomycin were inhibited to a similar or greater extent than their susceptible counterparts.

### Commensal treatments provide robust inhibition of MRSA over time

Having observed strong inhibition of JE2 by commensal species over 24h, we next asked whether inhibition was sustained over longer timescales. Our rationale here was that prolonged exposure to inhibitory commensal communities may result in complete eradication of JE2. Alternatively, if longer-term population dynamics differ from what we observed over a single growth cycle, or if JE2 adapts via genetic changes that overcome inhibition, then inhibition might not last. We investigated this by performing serial passage experiments over several days, co-passaging JE2 with commensal species for five growth cycles. At each transfer, we resuspended the entire community (including JE2 and commensal species) and transferred an aliquot to fresh media (see Methods). With this protocol, JE2 recovered high abundance by the second transfer in most treatment groups (**Figure 3A**), overcoming the inhibition we had observed previously over 24h. The only treatments where inhibition was sustained over all replicates were with treatments containing the commensal species of *S. aureus*. Consequently, final abundance of JE2 varied among different commensal-species treatments (effect of treatment in one-way ANOVA: F_6,35_ = 23.82, *p* < 0.001).

**Figure 3.**
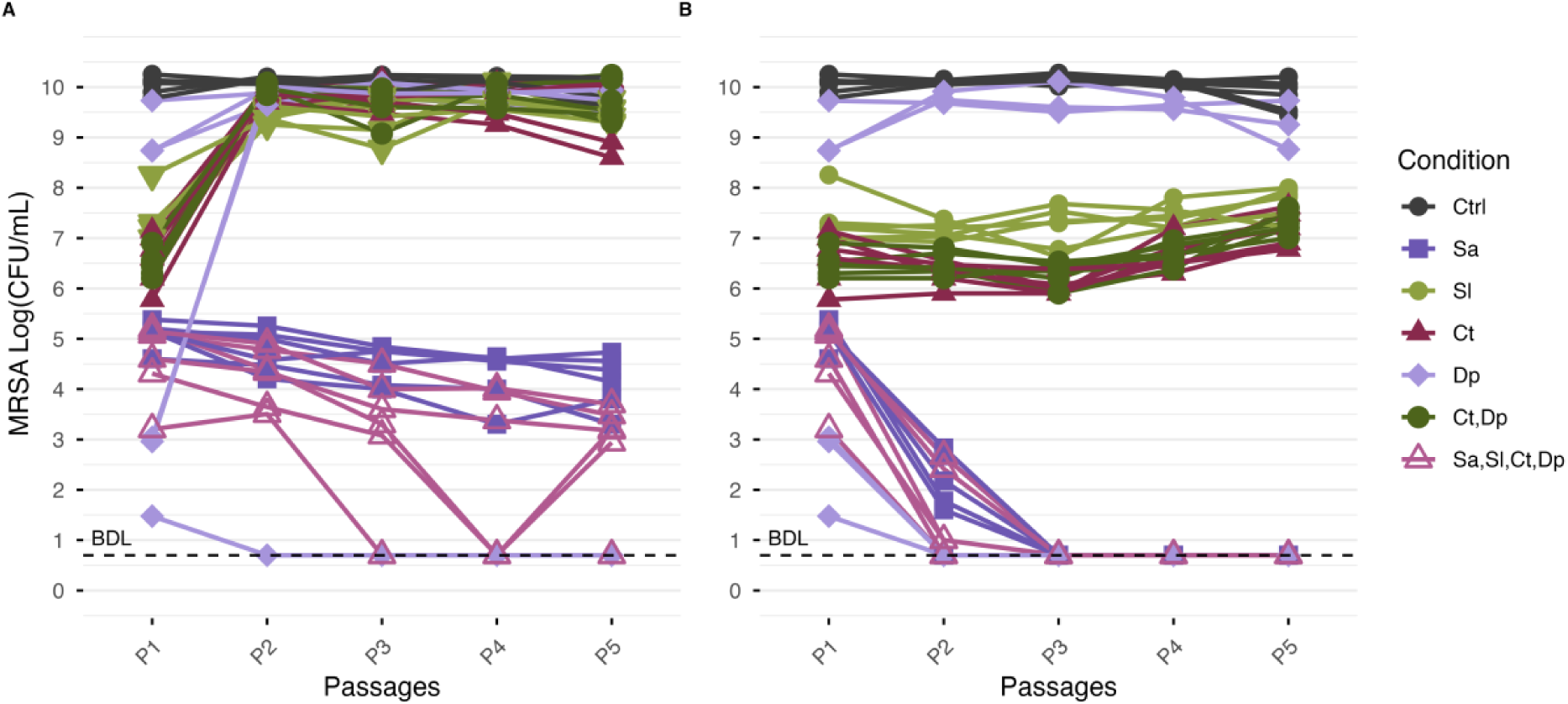
Commensal communities provide robust JE2 inhibition overtime. Growth of JE2 in communities of commensal species passaged five times on fresh medium (**A**) or fresh lawns of commensals (**B**). In both experiments, final JE2 abundance at the endpoint (P5) differed significantly among treatments (see main text); in **A**, treatments Sa and Sa,Sl,Ct,Dp differed significantly from the control, whereas in **B**, treatments Sa, Dp, and Sa,Sl,Ct,Dp differed significantly from the control (Dunnett’s test comparing each condition to Ctrl). Six independent replicates per condition. Ctrl: no commensal, Sa: *S. aureus*, Sl: *S. lugdunensis*, Ct: *C. tuberculostearicum*, Dp: *D. pigrum*.

We suspected the rapid recovery of JE2 during serial passage was due to its relatively high intrinsic growth rate compared to commensal species. This may allow JE2 to quickly outgrow other species after the community is transferred to fresh medium, unlike in our earlier colonisation assays where commensal species were first cultivated for 48h to establish a resident community. To account for this, we used a modified serial transfer protocol, where we transferred the culture atop a fresh lawn of commensals pre-grown for 48h, effectively reseeding a stable commensal community at each passage. In these conditions, final JE2 abundance again varied depending on which commensal-species treatment we applied (effect of treatment in one-way ANOVA: F_6,35_ = 22.73, p < 0.001). However, inhibition in multiple commensal-species treatments was sustained over time, including those with *S. lugdunensis*, *C. tuberculostearicum*, and the combination of *C. tuberculostearicum* and *D. pigrum*. Both treatment groups containing *S. aureus* pushed JE2 beneath the detection limit by the second and third passage (**Figure 3B**). In summary, the commensal species provided robust and sustained colonisation resistance under growth conditions that limited JE2 overgrowth during serial passaging.

### Partial adaptation of MRSA to inhibition by commensals

Pathobionts that reside in microbiomes are exposed to commensals for extended periods of time, which may result in adaptation. We next used a modified protocol to further investigate possible evolutionary adaptation of JE2 to overcome inhibition by commensals, which may also provide information about mechanisms by which commensals inhibit MRSA. Some of our serial transfer protocols in the above experiment caused complete eradication of JE2, so we aimed here to maintain JE2 population growth over longer timescales during serial passaging. To do this, we modified the protocol used above by plating the communities on MRSA-selective agar between transfer cycles and using recovered JE2 CFUs to inoculate a fresh lawn of commensals. We repeated this procedure for 20 transfer cycles, for six separate commensal treatments in five independent replicates each. We then isolated a single colony from each replicate population and tested for inhibition by commensals, compared with ancestral JE2. This showed several isolates that had been passaged with *C. tuberculostearicum* alone or *C. tuberculostearicum*+*D. pigrum* were less strongly inhibited by those commensal treatments compared to the ancestral strain (**Figure 4A**). Despite this, the same isolates were still strongly inhibited by these treatments compared to their growth in the absence of commensals (**Figure 4A**), indicating any adaptation to overcome commensal inhibition was partial. Isolates passaged with *C. tuberculostearicum*, *D. pigrum* or their combination also exhibited increased doubling time compared to the ancestral strain (**Figure S4**). Passaging with *S. aureus*, *S. lugdunensis*, *D. pigrum* and the four-species combination did not result in increased resistance to inhibition. To test for further potential pleiotropic effects of commensal adaptation, we assessed antibiotic susceptibility of adapted strains and found no major changes, aside from a modest increase in teicoplanin susceptibility in strains adapted to *C. tuberculostearicum* alone or in combination with *D. pigrum* (**Table S6**).

**Figure 4.**
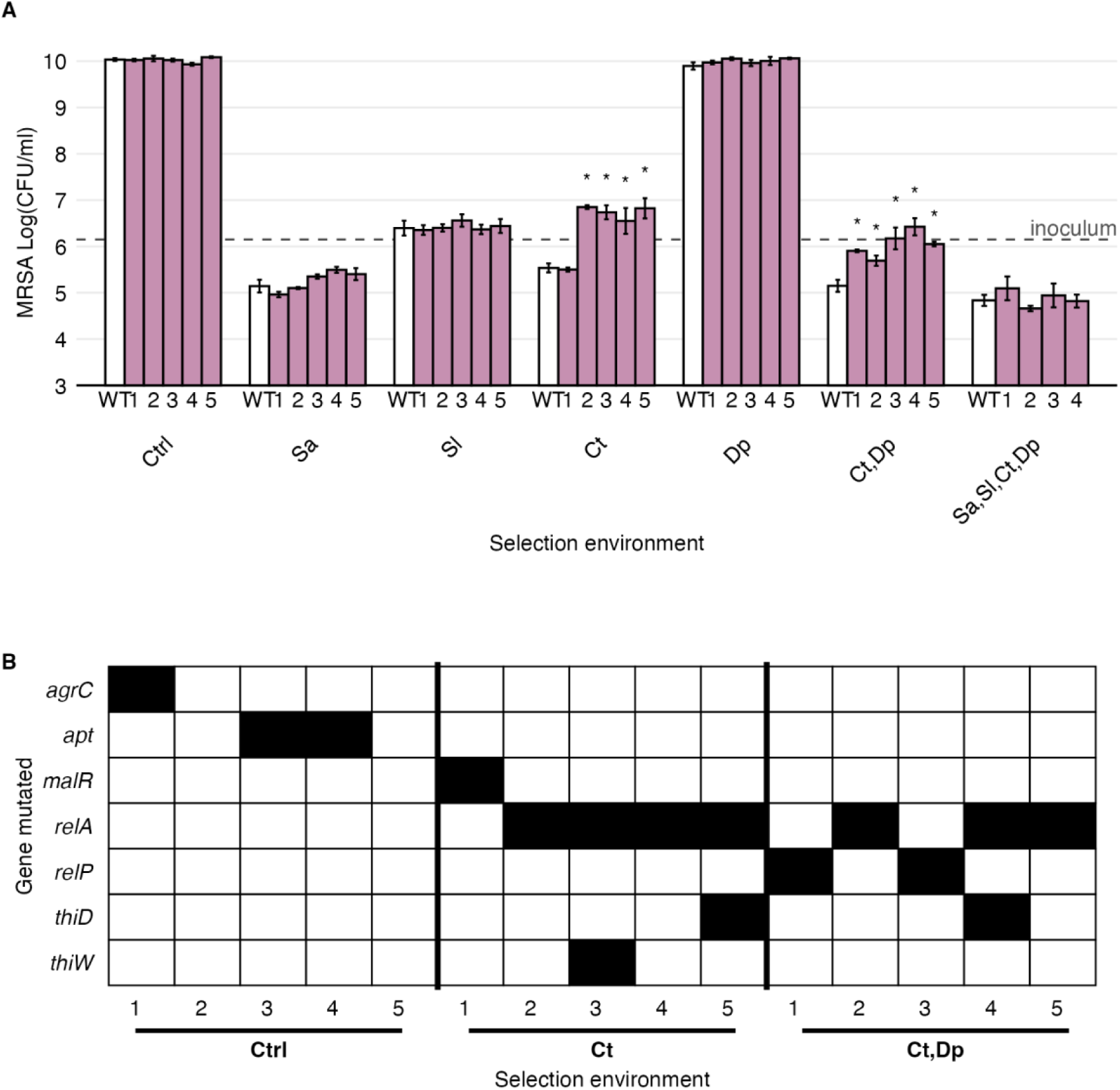
Adaptation of MRSA strain JE2 to commensal species from the human nose. **A.** Growth of commensal-passaged JE2 (JE2_1, JE2_2, JE2_3, JE2_4 and JE2_5), co-cultured with combinations of commensal species, measured as log_10_(CFU/mL). Growth is tested in the same conditions used for passaging, with ancestral JE2 for comparison. Bars represent mean ± SE. Statistical comparisons were performed using Dunnett’s test within each condition. \**p* ≤ 0.05. Sa: *S. aureus*, Se: *S. lugdunensis*, Ct: *C. tuberculostearicum*, Dp: *D. pigrum*. **B.** Genetic changes identified in JE2 isolates passaged alone (Ctrl), with *C. tuberculostearicum* (Ct), or with *C. tuberculostearicum* and *D. pigrum* (Ct,Dp). Rows represent mutated genes and columns represent independently passaged isolates (JE2_1–JE2_5 in each treatment). Black squares indicate presence of a mutation in the corresponding gene and isolate. Further details of individual mutations are given in **Table S7**.

To investigate the genetic basis of adaptation to *C. tuberculostearicum* alone or in combination with *D. pigrum*, we sequenced isolates from serially passaged JE2 populations. This revealed some loci mutated in multiple independently evolved isolates (**Figure 4B; Table S7**) and the same genes were more likely to be mutated in isolates from the same rather than different treatments (Permutation test: F_2, 12_ = 2.012, p = 0.022). For example, isolates passaged with *C. tuberculostearicum* acquired mutations in *relA* (7 out of 10 isolates), a central mediator of the stringent stress response, and mutations in genes linked with thiamine transport and biosynthesis (*thiD* and *thiW*). Other mutated loci included *relP*, a small alarmone synthetase associated with cell envelope stress responses (two out of five isolates in the *C. tuberculostearicum* and *D. pigrum* co-treatment). To further investigate the role of these genes (*relA*, *relP*, *thiD*, and *thiW*) in the interaction between *S. aureus* and *C. tuberculostearicum* with or without *D. pigrum*, we tested corresponding loss-of-function mutants from the Nebraska Transposon Mutant Library (Fey et al., 2013) in colonisation assays. Here, the effects of commensal treatments varied among tested strains (Strain × Condition interaction in two-way ANOVA: F_12, 63_ = 5.344, *p* < 0.001; **Figure S5**), but in most cases mutants responded similarly compared with the wild type. Together, these results suggest the observed partial adaptation to *C. tuberculostearicum*, alone or in combination with *D. pigrum*, involves genetic changes in stress-response and metabolic pathways, but that the net effects of such changes on susceptibility to inhibition may be specific to particular point mutations (we tested loss-of-function mutants in **Figure S5**, which may have different effects compared to mutations observed in passaged clones, **Table S7**) or combinations of mutations.

## Discussion

We sought to determine which bacterial species from the human nasal microbiota were most effective at inhibiting colonisation by MRSA, and to assess whether resistance to last-line antibiotics influenced this interaction. We assembled all possible combinations of highly prevalent nasal commensals in a fully factorial colonisation assay, allowing us to disentangle the effects of community complexity from those of specific taxa and to identify keystone species driving MRSA suppression. We found that Staphylococcal species inhibited growth of MRSA strain JE2. Similar levels of inhibition were achieved by combining *C. tuberculostearicum* and *D. pigrum*, with evidence that *D. pigrum* promotes growth of *Corynebacterium* species. Inhibition of MRSA by commensals was sustained over time and intraspecific competition effectively removed MRSA from the community. Importantly, nasal commensals were equally efficient at inhibiting MRSA isolates resistant to last-line antibiotics, highlighting the potential of commensal-based strategies to reduce the burden of antibiotic-resistant infections.

Our finding that intraspecific competition efficiently inhibited MRSA (**Figure 1**) is consistent with previous findings (Brülisauer et al., 2026), and indicates individual *S. aureus* carrier status may be a key determinant of susceptibility to MRSA colonisation. Consistent with this being relevant in nature, earlier work has shown that a single *S. aureus* strain colonises the nasal microbiome in 90% of carriers, and nasal carriage is highly dynamic, with transient co-carriage and strain replacement (Votintseva et al., 2014). This fits with the general expectation that closely related taxa may sometimes, but not always, compete relatively strongly, for example due to relatively high niche overlap (Godoy et al., 2014; Narwani et al., 2013; Webb et al., 2002). This is also evidenced by recent studies using synthetic communities modelling the human gut microbiome, showing resistance to pathogen colonisation is strengthened by the presence of closely related species (Spragge et al., 2023). Our results advance on these findings by providing direct experimental evidence that competition with other Staphylococci is sufficient to drive MRSA exclusion in a controlled colonisation model. This also aligns with evidence that some *S. lugdunensis* strains can inhibit *S. aureus* via production of the antibiotic lugdunin, while their persistence in the nasal cavity may be supported by other commensals (Rosenstein et al., 2024; Zipperer et al., 2016).

We observed strong MRSA inhibition in several treatments with non-staphylococcal commensal species, particularly combinations of *Corynebacterium* species and *D. pigrum*. That this type of inhibition may influence *S. aureus* colonisation in nature is supported by previous studies reporting negative associations between *D. pigrum*/*Corynebacterium* spp. and *S. aureus* in the nasal microbiome (Brugger et al., 2020; Liu et al., 2015), and reduced *S. aureus* colonisation when *D. pigrum* and *Corynebacterium pseudodiphteriticum*/*propinquum* co-occurred (Ingham et al., 2025; Liu et al., 2026). Notably, although direct inhibition of *S. aureus* by *D. pigrum* has been reported in agar-based assays (Brugger et al., 2020), *D. pigrum* alone did not consistently inhibit MRSA in our experiments, suggesting context-dependent effects. *Corynebacterium* species have been considered in the context of new antistaphylococcal interventions (Shamsuzzaman et al., 2023; Tepekule et al., 2025; Uehara et al., 2000). In vitro, *D. pigrum* and *Corynebacterium pseudodiphtheriticum* have also been shown to synergistically suppress *S. pneumoniae* (Brugger et al., 2020; Cisneros et al., 2025). Because *D. pigrum* exhibits multiple auxotrophies, it likely relies on both the host and co-colonizing microbes for essential nutrients (Brugger et al., 2020). Our results provide novel information about such interactions by demonstrating *D. pigrum* supports *Corynebacterium* growth, associated with enhanced MRSA inhibition. Furthermore, we found these interactions apply across multiple strains and species of the relevant genera, but with some degree of specificity: in supernatant assays, *D. pigrum* supported growth of *C. tuberculostearicum* and *C. accolens*, but not *C. propinquum*. A possible mechanism for this interaction may involve production of metabolites such as lactate, which can serve as a carbon source for some *Corynebacterium* spp. (Brugger et al., 2020; Stansen et al., 2005). In addition, *D. pigrum* expresses esterases and lipases capable of degrading exogenous lipids (Lissin et al., 2025; Parte et al., 2020), potentially releasing free fatty acids required for growth of lipophilic species. The specificity we observed among *D. pigrum*-*Corynebacterium* combinations is consistent with this: *C. tuberculostearicum* and *C. accolens* are both lipophilic, *C. propinquum* is not. The relatively strong inhibition we observed for some combination treatments is also in line with community-dependent effects observed in other host-associated microbiotas, where synthetic consortia can provide stronger colonisation resistance than individual strains alone (Wei et al., 2023). Similarly, a recent study showed a combination of two protective commensals broadened their inhibitory spectrum, suppressing a wider range of Enterobacterales than either strain individually (Wende et al., 2025). Thus, our results provide new information about the minimum community complexity required for maximal suppression of *S. aureus*.

The importance of nutrients, including fatty acids, in determining how commensals inhibit *S. aureus* is further supported by our finding that supplementation with an additional source of fatty acids benefited *Corynebacterium* species and led to increased inhibition of MRSA. Lipophilic *Corynebacterium* species like *C. tuberculostearicum* and *C. accolens* lack a complete fatty acid biosynthesis pathway and therefore depend on exogenous lipids for growth (Funke et al., 1997). This requirement likely drives their adaptation to lipid-rich environments, including sebaceous skin sites and the nasal cavity. It also helps to explain both the effect of tween supplementation in our experiments and potentially, as discussed above, the interactions between these *Corynebacterium* species and *D. pigrum*. More generally, nutrient limitation is relevant in the nasal passage (Krismer et al., 2014). Recent work indicates resource limitation and exchange with other species may play key roles in colonisation: some *S. aureus* strains are auxotrophic for tyrosine and rely on cross-feeding to establish colonisation (Bilici et al., 2025; Camus et al., 2025). Results from other types of microbiomes also support resource scarcity influencing the types of interactions among bacteria (Sulaiman et al., 2025). Thus, our results provide new insights into how environmental context, in this case, nutrient composition via fatty acid supplementation, can influence the structure and outcomes of microbial interactions.

Our results indicate that inhibition of MRSA by commensals is not decreased by resistance to anti-MRSA antibiotics. This is encouraging for the potential of commensals as probiotics against antibiotic-resistant pathogens, as such interventions are especially needed in contexts where antibiotics are no longer effective (Ghosh et al., 2019; Tepekule et al., 2025). The resistant strains we analysed carried representative mutations in genes involved in cell biosynthesis, transcription and DNA repair, and metabolism and nutrient acquisition. As for other resistant isolates of *S. aureus* and other species, they also had variable growth costs in the absence of antibiotics (Andersson & Hughes, 2010; Andersson & Levin, 1999). We also included clinical isolates. Nevertheless, we do not rule out other possible effects of resistance on colonisation and responses of MRSA to treatment: there is growing evidence that antibiotic resistance can affect ecological interactions and vice versa (Coll et al., 2025; Connor et al., 2023; Koch et al., 2014; Lindon et al., 2024; Pearl Mizrahi et al., 2023; Sulaiman et al., 2025; Vega et al., 2013). For instance, antibiotic resistance altered the ability of *Pseudomonas aeruginosa* to invade respiratory commensal communities in a species-and antibiotic-specific manner, resulting in either enhanced or reduced invasion (Lindon et al., 2024). Antibiotic resistance may affect other phenotypes, such as responses to antimicrobial molecules produced by the host mucosa as part of the innate immune system, and by resident bacteria that secrete bacteriocins (Cole et al., 2002; Fait et al., 2023; Iwase et al., 2010; Krismer et al., 2014; Zipperer et al., 2016). Indeed, our assay has limitations in that it is a simplified *in vitro* model that does not capture all traits of *in vivo* colonisation, such as host factors, spatial structure, and fluid flux, which may influence interaction outcomes. Thus, while our findings are promising in that commensal inhibition was robust to resistance against major anti-MRSA agents, testing for impacts on other relevant phenotypes before any downstream application will be informative.

Finally, while adaptation to abiotic stresses like antibiotics is well documented, bacteria can also evolve in response to biotic pressures imposed by other species (Sulaiman et al., 2025; Tognon et al., 2017). In our co-culture passaging experiments, MRSA adapted to inhibition by only some commensal species and did not fully overcome suppression. The JE2 strain partially adapted to inhibition by *C. tuberculostearicum*, both alone and in combination with *D. pigrum*, through mutations affecting metabolism, stress response, and the cell envelope. *S. aureus* adaptation to *C. pseudodiphtheriticum* inhibition was also associated with mutations affecting metabolic and cell envelope pathways, as well as quorum sensing (Hardy et al., 2019). Our findings are also consistent with prior observations of metabolic niche shift of the pathobiont *Clostridioides difficile* during prolonged co-culture with the gut commensal *Phocaeicola vulgatus*, and lipopolysaccharide mutations in *Pseudomonas aeruginosa* during co-culture with *S. aureus* (Sulaiman et al., 2025; Tognon et al., 2017). Together, these results indicate only limited capacity for MRSA to overcome commensal-mediated inhibition over the timescales and treatments tested here.

In conclusion, we identified commensal species and combinations that inhibited MRSA colonisation in an in vitro model. The relevance of several of our results, such as inhibition by *Corynebacterium*-*D. pigrum* combinations and the observed patterns of specificity among different strain combinations, is supported by past work on associations between taxonomic composition of the nasal microbiota and colonisation by *S. aureus* and in vitro tests in other set-ups. Crucially, in the context of potential future probiotic application or other microbiota-targeted interventions, these effects were consistent across strains with resistance to key anti-MRSA antibiotics, including clinical isolates.

## Methods

### Strains and growth conditions

Bacterial strains are listed in **Table S1**. We obtained commensal strains from the University Hospital Zürich. *Dolosigranulum pigrum* isolates were grown on Columbia agar supplemented with 5% sheep blood (BD Difco^TM^) at 34°C. Other commensal strains were grown on brain heart infusion agar (BHIA), a nutrient-rich medium that supports growth of fastidious organisms, supplemented with 1% Tween 80 at 34°C. For MRSA strains with additional resistance to last-line antibiotics, we included experimentally adapted isolates we generated and characterized for this project (Teicoplanin and Daptomycin; see below), experimentally adapted isolates from a previous work (Vancomycin; Fait et al., 2022), and pairs of clinical isolates from a previous work (Vancomycin; Howden et al., 2006). Experiments were conducted at 34°C, reflecting the average temperature in the nasopharynx, as *S. aureus* displays distinct transcriptional and proteomic profiles at this temperature compared to 37°C (Bastock et al., 2021; Keck et al., 2000). The Nebraska Transposon Mutant Library (NTML) was obtained from BEI resources and grown in BHI broth supplemented with 5mg/L erythromycin. Antibiotic stock solutions were prepared following manufacturer recommendations.

### Colonisation assay

To test the ability of bacterial commensal species from the human nose to inhibit growth of MRSA, we performed experiments on semi-solid surfaces to reflect nasal growth conditions and preserve spatial structure. We used this assay as a model of early-stage colonisation. BHIA with or without 1% Tween 80 was added to the wells of 12-well microplates. Commensal species were assembled in communities by resuspending colonies in buffer to a final OD_600nm_=1, before adding 50μL to microplate wells and incubating at 34°C for 48h, with four biological replicates. We included this first culturing step, with commensals alone, to prevent overgrowth by JE2. MRSA strains were resuspended from colonies to OD_600nm_=0.01 and 50μL was added to the commensal lawns. A low MRSA inoculum was used to reflect the low density of incoming pathogens relative to established commensals and simulate a hypothetical colonisation scenario. Plates were incubated at 34°C for 24h. Bacteria in each well were then resuspended in 1mL saline, diluted and plated on MRSA selective media (CHROMagar^TM^). After incubation, we estimated MRSA abundance by counting colony-forming units (CFU). To assess the contribution of niche occupation to JE2 inhibition, we estimated commensal abundance under these conditions prior to MRSA inoculation in a separate experiment. This was performed as above, except that samples were plated on BHIA + 1% Tween 80, and an additional set of plates was incubated anaerobically to account for growth of the aerotolerant anaerobe *C. acnes*.

### Teicoplanin and daptomycin adaptation

To isolate JE2 mutants with elevated resistance against teicoplanin and daptomycin, we serially passaged replicate JE2 cultures with increasing concentrations of each antibiotic. We began by inoculating 10 sets of replicate cultures from overnight cultures of JE2 into microplates containing various concentrations of antibiotics in 0.5mg/L increments. Next, after 24h at 37°C without shaking, we recorded the highest concentration for each replicate set of cultures where bacterial growth was visible in the plate well. We then transferred an aliquot of this individual well to a microtiter plate containing a fresh gradient of antibiotic concentrations in a 1:1000 dilution ratio. We repeated this process for ten growth cycles for daptomycin and four growth cycles for teicoplanin, until all replicate cultures had reached MICs above clinical breakpoints (EUCAST, 2024; Daptomycin, MIC >1mg/L; Teicoplanin, MIC > 2mg/L). During serial passage, we regularly plated cultures on BHIA to visually check for contaminants. Growth media were supplemented with 50 mg/L calcium in daptomycin treated cultures, following EUCAST recommendations (EUCAST, 2024). At the end of serial passaging, we plated the replicate cultures on BHIA and picked a single colony from each. The antibiotic-adapted isolates were saved in glycerol at-75°C and characterised by determining their MICs and doubling time. We selected three vancomycin-adapted JE2-derived isolates from a previous study based on their phenotypic diversity, with VAN1 and VAN9 showing increased susceptibility to β-lactam antibiotics and VAN4 displayed decreased susceptibility (Fait et al., 2022).

### Generation times

To estimate growth cost associated with resistance (**Figure S3**) and adaptation to commensals (**Figure S4**), we diluted overnight cultures OD_600nm_=0.01 in BHI and incubated at 37°C in a microplate reader (Tecan) for 24h with OD_600nm_ measurements at 15min intervals. Each isolate was tested in three biological replicates. Doubling times were calculated from using the fit_easylinear function from the growthrates R package (v0.8.5, Hall et al., 2014). Doubling times were calculated as ln(2) divided by the maximum exponential growth rate and reported in minutes.

### Antibiotic susceptibility testing

Minimum inhibitory concentrations (MIC) for daptomycin-and teicoplanin-adapted JE2 and for commensal-adapted JE2 were determined using antibiotic strips (Biomérieux) according to instructions from the manufacturer.

### Whole genome sequencing and analysis

Sequencing was performed at the Microbial Genomics Division of the Institute of Medical Microbiology, University of Zürich. DNA extractions were performed using the Maxwell® RSC platform with the Maxwell® RSC Blood DNA Kit. Library preparations were performed using the QiaSeqFX kit (Qiagen) and sequencing was carried using Illumina NextSeq1000 platform PE150. Analysis of output reads was performed using CLC Genomics Workbench 24.0.2. Reads were aligned to the reference genome of *S. aureus* USA300_FPR3757 (NC_07793). Mutations were verified manually. Variants detected in JE2 WT were excluded in the adapted isolates.

### Serial passaging of JE2 with commensals

We conducted serial passaging experiments to test whether inhibition of MRSA by commensals was sustained over a longer timescale. BHI agar supplemented with 1% Tween 80 was added to the wells of a 12-well microplate. Commensal species were assembled in communities of final OD_600nm_=1. Wells were inoculated with 50uL of each community and incubated for 48h at 34°C. 50μL of JE2 diluted to OD_600nm_=0.1 was added to the commensal lawns in six independent replicates. Plates were incubated for 24h at 34°C. Wells were resuspended in 1mL saline. Communities were passaged by adding 50μL of the resuspended community atop fresh media and incubating 24h at 34°C. Communities were diluted and plated on MRSA-selective agar to monitor the abundance of JE2 in log_10_(CFU/mL). We carried this out for five replicate cultures for a total of 20 growth cycles. We used two further variations of this protocol. First, to maintain substantial commensal abundance during serial passage, we transferred 50μL of the resuspended community to a pre-grown lawn of commensals, instead of fresh media, at each transfer. Second, to maintain JE2 abundance in testing for evolutionary adaptation to commensal treatments, we isolated and re-introduced JE2 at each transfer, by plating aliquots on MRSA-selective agar, pooling and resuspending the MRSA colonies, and diluting to OD_600nm_=0.01 to inoculate on fresh lawns of commensals. JE2 was passaged in these conditions for 20 cycles, in five independent replicates for each commensal condition. The final communities were plated on MRSA-selective agar, and three colonies were picked per replicate per condition and saved in glycerol stocks at-75°C. One colony per replicate per condition was used in colonisation assays to compare the inhibition of passaged-JE2 and ancestral JE2 by commensals in sympatric conditions (in the commensal conditions used for passaging).

### Supernatant assay

To determine whether growth of the commensal species in our colonisation-assay set-up altered the abiotic environment in a way that affected growth of other species, we collected supernatant from cultures of different commensals. First, we cultivated single-species lawns of all commensals in the same conditions as during our colonisation assays in three replicates. We then resuspended each culture in saline and filtered (0.22 μm) to obtain a sterile supernatant. As a control treatment, we carried out the same procedure but with empty wells that had not been inoculated with commensals. To monitor growth of other bacteria in each supernatant, we then inoculated microplate wells, each filled with 50μl supernatant or control-supernatant, with the relevant test species from resuspended colonies in saline to achieve a final OD_600nm_= 0.01 in a final volume of 100μL. We incubated the microplates in a shaking incubator at 120rpm for 24h at 34°C before recording the final OD_600nm_ (**Figure 1D**, **Table S3**).

### Statistics

All statistical analysis was done in R 4.5.2 (R Core Team, 2025). In cases where we compared multiple treatments to a common control treatment, we used Dunnett’s Test (Dunnett, 1955). In other cases where multiple tests were made, we used Holm-Bonferroni (Holm, 1979) correction. To test the effects of individual commensal species on final JE2 abundance, we used a linear model with final JE2 abundance as the response variable and presence/absence of each species across treatments as factors; we performed stepwise reduction of this model using the stepAIC function from the MASS package (Venables & Ripley, 2002) using the penalty term *k = log(n),* where n is equal to sample size (i.e., using the Bayesian Information Criterion, rather than the Akaike Information Criterion), to compare models. To test whether coculture with *C. tuberculostaricum* and *D. pigrum* led to mutations in different genes, we used permutational analysis of variance (Anderson, 2001). Genotypes were scored with 1 (mutated) and 0 (not mutated) for each gene, before the square root of the Euclidian distance was calculated. F-tests comparing observed values to random permutations of the raw data were conducted using the function adonis2 in the R package vegan (version 2.7-2).

## Data availability

### Code availability

All code is available at ADDLINK.

## Acknowledgements

This work was supported by the Swiss National Science Foundation (A.H., project n° 310030_192428; S.D.B., project n° 211422). We thank Dr. Jeruscha Baum and Dr. Willy Isao Staiger for providing commensal isolates, and the group of Prof. Silvio D. Brugger for valuable feedback. We also thank Prof. Timothy P. Stinear and Dr. Ian R. Monk for providing VISA clinical isolates.

## Author contributions

A.F., and A.R.H. conceived the study and designed the experiments. A.F., A.R.H. S.D.B. and L.B. developed the methodology. A.F. and V.S. performed the experiments. A.F. and D.C.A. undertook bioinformatic and statistical analyses and generated the figures. A.R.H. secured funding. A.F. and A.R.H. wrote the original manuscript, and all authors read, reviewed and approved the manuscript.

## Materials & correspondence

Correspondence and material requests should be addressed to A.F.

## References

Adolf, L. A., & Heilbronner, S. (2022). Nutritional Interactions between Bacterial Species Colonising the Human Nasal Cavity: Current Knowledge and Future Prospects. Metabolites, 12(6), 489. 10.3390/metabo12060489

Anderson, M. J. (2001). A new method for non-parametric multivariate analysis of variance. Austral Ecology, 26(1), 32–46. 10.1111/j.1442-9993.2001.01070.pp.x

Bastock, R. A., Marino, E. C., Wiemels, R. E., Holzschu, D. L., Keogh, R. A., Zapf, R. L., Murphy, E. R., & Carroll, R. K. (2021). Staphylococcus aureus Responds to Physiologically Relevant Temperature Changes by Altering Its Global Transcript and Protein Profile. mSphere, 6(2), e01303–20. 10.1128/mSphere.01303-20

Bilici, K., Gerlach, D., Camus, L., & Heilbronner, S. (2025). Competitive fitness of Staphylococcus aureus against nasal commensals depends on biotin biosynthesis and acquisition. The ISME Journal, 19(1), wraf248. 10.1093/ismejo/wraf248

Boyle-Vavra, S., Yin, S., Jo, D. S., Montgomery, C. P., & Daum, R. S. (2013). VraT/YvqF Is Required for Methicillin Resistance and Activation of the VraSR Regulon in Staphylococcus aureus. Antimicrobial Agents and Chemotherapy, 57(1), 83–95. 10.1128/AAC.01651-12

Brugger, S. D., Eslami, S. M., Pettigrew, M. M., Escapa, I. F., Henke, M. T., Kong, Y., & Lemon, K. P. (2020). Dolosigranulum pigrum Cooperation and Competition in Human Nasal Microbiota. mSphere, 5(5), e00852–20. 10.1128/mSphere.00852-20

Brülisauer, L., Boumasmoud, M., Fait, A., Brugger, S. D., & Hall, A. R. (2026). Individual bacterial taxa drive colonisation resistance to methicillin-resistant Staphylococcus aureus in human nasal microbiome samples (p. 2026.01.07.698121). bioRxiv. 10.64898/2026.01.07.698121

Camus, L., Franz, J., Gerlach, D., Lange, A., Power, J. J., Navarro-Díaz, M., Rapp, J., Otto, M., Klages, L. J., Starostin, V., Mößner, J., Bilici, K., Ham, S., Kalinowski, J., Angenent, L. T., Link, H., & Heilbronner, S. (2025). Tyrosine availability shapes Staphylococcus aureus nasal colonization and interactions with commensal communities (p. 2025.05.06.651429). bioRxiv. 10.1101/2025.05.06.651429

Cisneros, M., Blanco-Fuertes, M., Lluansí, A., Brotons, P., Henares, D., Pérez-Argüello, A., González-Comino, G., Ciruela, P., Mira, A., & Muñoz-Almagro, C. (2025). Synergistic inhibition of pneumococcal growth by Dolosigranulum pigrum and Corynebacterium pseudodiphtheriticum: Insights into nasopharyngeal microbial interactions. Microbiology Spectrum, 13(7), e00138–25. 10.1128/spectrum.00138-25

Clarridge, J. E., Harrington, A. T., Roberts, M. C., Soge, O. O., & Maquelin, K. (2013). Impact of strain typing methods on assessment of relationship between paired nares and wound isolates of methicillin-resistant Staphylococcus aureus. Journal of Clinical Microbiology, 51(1), 224–231. 10.1128/JCM.02423-12

Cole, A. M., Liao, H.-I., Stuchlik, O., Tilan, J., Pohl, J., & Ganz, T. (2002). Cationic Polypeptides Are Required for Antibacterial Activity of Human Airway Fluid. The Journal of Immunology, 169(12), 6985–6991. 10.4049/jimmunol.169.12.6985

Coll, F., Blane, B., Bellis, K. L., Matuszewska, M., Wonfor, T., Jamrozy, D., Toleman, M. S., Geoghegan, J. A., Parkhill, J., Massey, R. C., Peacock, S. J., & Harrison, E. M. (2025). The mutational landscape of Staphylococcus aureus during colonisation. Nature Communications, 16(1), 302. 10.1038/s41467-024-55186-x

Connor, C. H., Zucoloto, A. Z., Munnoch, J. T., Yu, I.-L., Corander, J., Hoskisson, P. A., McDonald, B., & McNally, A. (2023). Multidrug-resistant E. coli encoding high genetic diversity in carbohydrate metabolism genes displace commensal E. coli from the intestinal tract. PLOS Biology, 21(10), e3002329. 10.1371/journal.pbio.3002329

Cui, L., Isii, T., Fukuda, M., Ochiai, T., Neoh, H., Camargo, I. L. B. da C., Watanabe, Y., Shoji, M., Hishinuma, T., & Hiramatsu, K. (2010). An RpoB Mutation Confers Dual Heteroresistance to Daptomycin and Vancomycin in Staphylococcus aureus. Antimicrobial Agents and Chemotherapy, 54(12), 5222–5233. 10.1128/AAC.00437-10

Dubrac, S., Bisicchia, P., Devine, K. M., & Msadek, T. (2008). A matter of life and death: Cell wall homeostasis and the WalKR (YycGF) essential signal transduction pathway. Molecular Microbiology, 70(6), 1307–1322. 10.1111/j.1365-2958.2008.06483.x

Dunnett, C. W. (1955). A Multiple Comparison Procedure for Comparing Several Treatments with a Control. Journal of the American Statistical Association, 50(272), 1096–1121. 10.1080/01621459.1955.10501294

Escapa, I. F., Chen, T., Huang, Y., Gajare, P., Dewhirst, F. E., & Lemon, K. P. (2018). New Insights into Human Nostril Microbiome from the Expanded Human Oral Microbiome Database (eHOMD): A Resource for the Microbiome of the Human Aerodigestive Tract. mSystems, 3(6), e00187–18. 10.1128/mSystems.00187-18

EUCAST. (2024). The European Committee on Antimicrobial Susceptibility Testing. Breakpoint tables for interpretation of MICs and zone diameters. Version 14.0, 2024. http://www.eucast.org.

Fait, A., Andersson, D. I., & Ingmer, H. (2023). Evolutionary history of Staphylococcus aureus influences antibiotic resistance evolution. Current Biology, 33(16), 3389–3397.e5. 10.1016/j.cub.2023.06.082

Fait, A., Seif, Y., Mikkelsen, K., Poudel, S., Wells, J. M., Palsson, B. O., & Ingmer, H. (2022). Adaptive laboratory evolution and independent component analysis disentangle complex vancomycin adaptation trajectories. Proceedings of the National Academy of Sciences, 119(30), e2118262119. 10.1073/pnas.2118262119

Fait, A., Silva, S. F., Abrahamsson, J. Å. H., & Ingmer, H. (2024). *Staphylococcus aureus* response and adaptation to vancomycin. In R. K. Poole & D. J. Kelly (Eds.), Advances in Microbial Physiology (Vol. 85, pp. 201–258). Academic Press. 10.1016/bs.ampbs.2024.04.006

Fey, P. D., Endres, J. L., Yajjala, V. K., Widhelm, T. J., Boissy, R. J., Bose, J. L., & Bayles, K. W. (2013). A Genetic Resource for Rapid and Comprehensive Phenotype Screening of Nonessential Staphylococcus aureus Genes. mBio, 4(1), e00537–12. 10.1128/mBio.00537-12

Freeman, C. D., Hansen, T., Urbauer, R., Wilkinson, B. J., Singh, V. K., & Hines, K. M. (2024). Defective pgsA contributes to increased membrane fluidity and cell wall thickening in Staphylococcus aureus with high-level daptomycin resistance. mSphere, 9(6), e00115–24. 10.1128/msphere.00115-24

Funke, G., von Graevenitz, A., Clarridge, J. E., & Bernard, K. A. (1997). Clinical microbiology of coryneform bacteria. Clinical Microbiology Reviews, 10(1), 125–159. 10.1128/cmr.10.1.125

Ghosh, C., Sarkar, P., Issa, R., & Haldar, J. (2019). Alternatives to Conventional Antibiotics in the Era of Antimicrobial Resistance. Trends in Microbiology, 27(4), 323–338. 10.1016/j.tim.2018.12.010

Godoy, O., Kraft, N. J. B., & Levine, J. M. (2014). Phylogenetic relatedness and the determinants of competitive outcomes. Ecology Letters, 17(7), 836–844. 10.1111/ele.12289

Hall, B. G., Acar, H., Nandipati, A., & Barlow, M. (2014). Growth rates made easy. Molecular Biology and Evolution, 31(1), 232–238. 10.1093/molbev/mst187

Hardy, B. L., Bansal, G., Hewlett, K. H., Arora, A., Schaffer, S. D., Kamau, E., Bennett, J. W., & Merrell, D. S. (2020). Antimicrobial Activity of Clinically Isolated Bacterial Species Against Staphylococcus aureus. Frontiers in Microbiology, 10. https://www.frontiersin.org/articles/10.3389/fmicb.2019.02977

Hardy, B. L., Dickey, S. W., Plaut, R. D., Riggins, D. P., Stibitz, S., Otto, M., & Merrell, D. S. (2019). Corynebacterium pseudodiphtheriticum Exploits Staphylococcus aureus Virulence Components in a Novel Polymicrobial Defense Strategy. mBio, 10(1), e02491–18. 10.1128/mBio.02491-18

Hartman, B. J., & Tomasz, A. (1984). Low-affinity penicillin-binding protein associated with beta-lactam resistance in Staphylococcus aureus. Journal of Bacteriology, 158(2), 513–516.

Holm, S. (1979). A Simple Sequentially Rejective Multiple Test Procedure. Scandinavian Journal of Statistics, 6(2), 65–70.

Howden, B. P., Johnson, P. D. R., Ward, P. B., Stinear, T. P., & Davies, J. K. (2006). Isolates with Low-Level Vancomycin Resistance Associated with Persistent Methicillin-Resistant Staphylococcus aureus Bacteremia. Antimicrobial Agents and Chemotherapy, 50(9), 3039–3047. 10.1128/AAC.00422-06

Howden, B. P., McEvoy, C. R. E., Allen, D. L., Chua, K., Gao, W., Harrison, P. F., Bell, J., Coombs, G., Bennett-Wood, V., Porter, J. L., Robins-Browne, R., Davies, J. K., Seemann, T., & Stinear, T. P. (2011). Evolution of Multidrug Resistance during Staphylococcus aureus Infection Involves Mutation of the Essential Two Component Regulator WalKR. PLoS Pathogens, 7(11). 10.1371/journal.ppat.1002359

Ingham, A. C., Ng, D. Y. K., Iversen, S., Liu, C. M., Dinh, K. M., Holtfreter, S., Edslev, S. M., Johannesen, T. B., Rendboe, A. K., Christiansen, M. T., Ng, K. L., Skov, R., Samietz, S., Radke, D., Weiss, S., Völker, U., Bröker, B. M., Erikstrup, L. T., Erikstrup, C.,…Stegger, M. (2025). Staphylococci in high resolution: Capturing diversity within the human nasal microbiota. Cell Reports, 44(6). 10.1016/j.celrep.2025.115854

Iwase, T., Uehara, Y., Shinji, H., Tajima, A., Seo, H., Takada, K., Agata, T., & Mizunoe, Y. (2010). Staphylococcus epidermidis Esp inhibits Staphylococcus aureus biofilm formation and nasal colonization. Nature, 465(7296), Article 7296. 10.1038/nature09074

Keck, T., Leiacker, R., Riechelmann, H., & Rettinger, G. (2000). Temperature Profile in the Nasal Cavity. The Laryngoscope, 110(4), 651–654. 10.1097/00005537-200004000-00021

Koch, G., Yepes, A., Förstner, K. U., Wermser, C., Stengel, S. T., Modamio, J., Ohlsen, K., Foster, K. R., & Lopez, D. (2014). Evolution of resistance to a last-resort antibiotic in Staphyloccocus aureus via bacterial competition. Cell, 158(5), 1060–1071. 10.1016/j.cell.2014.06.046

Krismer, B., Liebeke, M., Janek, D., Nega, M., Rautenberg, M., Hornig, G., Unger, C., Weidenmaier, C., Lalk, M., & Peschel, A. (2014). Nutrient Limitation Governs Staphylococcus aureus Metabolism and Niche Adaptation in the Human Nose. PLoS Pathogens, 10(1), e1003862. 10.1371/journal.ppat.1003862

Lindon, S., Shah, S., Gifford, D. R., Lood, C., Gomis Font, M. A., Kaur, D., Oliver, A., MacLean, R. C., & Wheatley, R. M. (2024). Antibiotic resistance alters the ability of Pseudomonas aeruginosa to invade bacteria from the respiratory microbiome. *Evolution Letters*, qrae030. 10.1093/evlett/qrae030

Lissin, A., Schober, I., Witte, J. F., Lüken, H., Podstawka, A., Koblitz, J., Bunk, B., Dawyndt, P., Vandamme, P., de Vos, P., Overmann, J., & Reimer, L. C. (2025). StrainInfo—The central database for linked microbial strain identifiers. Database, 2025, baaf059. 10.1093/database/baaf059

Liu, C. M., Erikstrup, L. T., Edslev, S. M., Park, D. E., Salazar, J. E., Aziz, M., Rendboe, A. K., Pham, T., Dinh, K. M., Roach, K., Onos, A., Sung, E., Weber, N. O., Andersen, P. S., Ullum, H., Skov, R., Hungate, B. A., Stegger, M., Erikstrup, C., & Price, L. B. (2026). Composition and dynamics of the adult nasal microbiome. Microbiome, 14(1), 38. 10.1186/s40168-025-02250-3

Liu, C. M., Price, L. B., Hungate, B. A., Abraham, A. G., Larsen, L. A., Christensen, K., Stegger, M., Skov, R., & Andersen, P. S. (2015a). Staphylococcus aureus and the ecology of the nasal microbiome. Science Advances, 1(5), e1400216. 10.1126/sciadv.1400216

Liu, C. M., Price, L. B., Hungate, B. A., Abraham, A. G., Larsen, L. A., Christensen, K., Stegger, M., Skov, R., & Andersen, P. S. (2015b). Staphylococcus aureus and the ecology of the nasal microbiome. Science Advances, 1(5), e1400216. 10.1126/sciadv.1400216

Mishra, N. N., McKinnell, J., Yeaman, M. R., Rubio, A., Nast, C. C., Chen, L., Kreiswirth, B. N., & Bayer, A. S. (2011). In Vitro Cross-Resistance to Daptomycin and Host Defense Cationic Antimicrobial Peptides in Clinical Methicillin-Resistant Staphylococcus aureus Isolates▿. Antimicrobial Agents and Chemotherapy, 55(9), 4012–4018. 10.1128/AAC.00223-11

Naghavi, M., Vollset, S. E., Ikuta, K. S., Swetschinski, L. R., Gray, A. P., Wool, E. E., Aguilar, G. R., Mestrovic, T., Smith, G., Han, C., Hsu, R. L., Chalek, J., Araki, D. T., Chung, E., Raggi, C., Hayoon, A. G., Weaver, N. D., Lindstedt, P. A., Smith, A. E.,…Murray, C. J. L. (2024). Global burden of bacterial antimicrobial resistance 1990–2021: A systematic analysis with forecasts to 2050. The Lancet, 404(10459), 1199–1226. 10.1016/S0140-6736(24)01867-1

Narwani, A., Alexandrou, M. A., Oakley, T. H., Carroll, I. T., & Cardinale, B. J. (2013). Experimental evidence that evolutionary relatedness does not affect the ecological mechanisms of coexistence in freshwater green algae. Ecology Letters, 16(11), 1373–1381. 10.1111/ele.12182

Pál, C., Papp, B., & Lázár, V. (2015). Collateral sensitivity of antibiotic-resistant microbes. Trends in Microbiology, 23(7), 401–407. 10.1016/j.tim.2015.02.009

Panwar, R. B., Sequeira, R. P., & Clarke, T. B. (2021). Microbiota-mediated protection against antibiotic-resistant pathogens. Genes & Immunity, 22(5), 255–267. 10.1038/s41435-021-00129-5

Parsons, J. B., Mourad, A., Conlon, B. P., Kielian, T., & Fowler, V. G. (2025). Methicillin-resistant and susceptible Staphylococcus aureus: Tolerance, immune evasion and treatment. Nature Reviews Microbiology, 1–19. 10.1038/s41579-025-01226-2

Parte, A. C., Sardà Carbasse, J., Meier-Kolthoff, J. P., Reimer, L. C., & Göker, M. (2020). List of Prokaryotic names with Standing in Nomenclature (LPSN) moves to the DSMZ. International Journal of Systematic and Evolutionary Microbiology, 70(11), 5607–5612. 10.1099/ijsem.0.004332

Paulander, W., Maisnier-Patin, S., & Andersson, D. I. (2009). The Fitness Cost of Streptomycin Resistance Depends on rpsL Mutation, Carbon Source and RpoS (σS). Genetics, 183(2), 539–546. 10.1534/genetics.109.106104

Pearl Mizrahi, S., Goyal, A., & Gore, J. (2023). Community interactions drive the evolution of antibiotic tolerance in bacteria. Proceedings of the National Academy of Sciences, 120(3), e2209043119. 10.1073/pnas.2209043119

Popowitch, E. B., Tran, T. H., Escapa, I. F., Bhatt, E., Sozat, A. K., Ahmed, N., Deming, C., Roberts, A. Q., Segre, J. A., Kong, H. H., Conlan, S., Lemon, K. P., & Kelly, M. S. (2024). Description of two novel Corynebacterium species isolated from human nasal passages and skin. bioRxiv, 2024.11.21.624533. 10.1101/2024.11.21.624533

Qi, Q., Preston, G. M., & MacLean, R. C. (2014). Linking System-Wide Impacts of RNA Polymerase Mutations to the Fitness Cost of Rifampin Resistance in Pseudomonas aeruginosa. mBio, 5(6), 10.1128/mbio.01562-14

Quinn, A. M., Bottery, M. J., Thompson, H., & Friman, V.-P. (2022). Resistance evolution can disrupt antibiotic exposure protection through competitive exclusion of the protective species. The ISME Journal, 16(10), 2433–2447. 10.1038/s41396-022-01285-w

Reynolds, P. E. (1989). Structure, biochemistry and mechanism of action of glycopeptide antibiotics. European Journal of Clinical Microbiology and Infectious Diseases, 8(11), 943–950. 10.1007/BF01967563

Rosenstein, R., Torres Salazar, B. O., Sauer, C., Heilbronner, S., Krismer, B., & Peschel, A. (2024). The Staphylococcus aureus-antagonizing human nasal commensal Staphylococcus lugdunensis depends on siderophore piracy. Microbiome, 12(1), 213. 10.1186/s40168-024-01913-x

Shamsuzzaman, M., Dahal, R. H., Kim, S., & Kim, J. (2023). Genome insight and probiotic potential of three novel species of the genus Corynebacterium. Frontiers in Microbiology, 14. 10.3389/fmicb.2023.1225282

Stansen, C., Uy, D., Delaunay, S., Eggeling, L., Goergen, J.-L., & Wendisch, V. F. (2005). Characterization of a Corynebacterium glutamicum Lactate Utilization Operon Induced during Temperature-Triggered Glutamate Production. Applied and Environmental Microbiology, 71(10), 5920–5928. 10.1128/AEM.71.10.5920-5928.2005

Sulaiman, J. E., Thompson, J., Cheung, P. L. K., Qian, Y., Mill, J., James, I., Im, H., Vivas, E. I., Simcox, J., & Venturelli, O. S. (2025). *Phocaeicola vulgatus* shapes the long-term growth dynamics and evolutionary adaptations of *Clostridioides difficile*. Cell Host & Microbe, 33(1), 42–58.e10. 10.1016/j.chom.2024.12.001

Tepekule, B., Barcik, W., Staiger, W. I., Bergadà-Pijuan, J., Scheier, T., Brülisauer, L., Hall, A. R., Günthard, H. F., Hilty, M., Kouyos, R. D., & Brugger, S. D. (2025). Computational and in vitro evaluation of probiotic treatments for nasal Staphylococcus aureus decolonization. Proceedings of the National Academy of Sciences, 122(7), e2412742122. 10.1073/pnas.2412742122

Thitiananpakorn, K., Aiba, Y., Tan, X.-E., Watanabe, S., Kiga, K., Sato’o, Y., Boonsiri, T., Li, F.-Y., Sasahara, T., Taki, Y., Azam, A. H., Zhang, Y., & Cui, L. (2020). Association of mprF mutations with cross-resistance to daptomycin and vancomycin in methicillin-resistant Staphylococcus aureus (MRSA). Scientific Reports, 10(1), Article 1. 10.1038/s41598-020-73108-x

Tognon, M., Köhler, T., Gdaniec, B. G., Hao, Y., Lam, J. S., Beaume, M., Luscher, A., Buckling, A., & van Delden, C. (2017). Co-evolution with Staphylococcus aureus leads to lipopolysaccharide alterations in Pseudomonas aeruginosa. The ISME Journal, 11(10), 2233–2243. 10.1038/ismej.2017.83

Uehara, Y., Nakama, H., Agematsu, K., Uchida, M., Kawakami, Y., Abdul Fattah, A. S. M., & Maruchi, N. (2000). Bacterial interference among nasal inhabitants: Eradication of *Staphylococcus aureus* from nasal cavities by artificial implantation of *Corynebacterium* sp. Journal of Hospital Infection, 44(2), 127–133. 10.1053/jhin.1999.0680

Vega, N. M., Allison, K. R., Samuels, A. N., Klempner, M. S., & Collins, J. J. (2013). Salmonella typhimurium intercepts Escherichia coli signaling to enhance antibiotic tolerance. Proceedings of the National Academy of Sciences, 110(35), 14420–14425. 10.1073/pnas.1308085110

von Eiff, C., Becker, K., Machka, K., Stammer, H., & Peters, G. (2001). Nasal Carriage as a Source of Staphylococcus aureus Bacteremia. New England Journal of Medicine, 344(1), 11–16. 10.1056/NEJM200101043440102

Votintseva, A. A., Miller, R. R., Fung, R., Knox, K., Godwin, H., Peto, T. E. A., Crook, D. W., Bowden, R., & Walker, A. S. (2014). Multiple-Strain Colonization in Nasal Carriers of Staphylococcus aureus. Journal of Clinical Microbiology, 52(4), 1192–1200. 10.1128/JCM.03254-13

Webb, C. O., Ackerly, D. D., McPeek, M. A., & Donoghue, M. J. (2002). Phylogenies and Community Ecology. Annual Review of Ecology, Evolution, and Systematics, 33(Volume 33, 2002), 475–505. 10.1146/annurev.ecolsys.33.010802.150448

Wei, M., Flowers, L., Knight, S. A. B., Zheng, Q., Murga-Garrido, S., Uberoi, A., Pan, J. T.-C., Walsh, J., Schroeder, E., Chu, E. W., Campbell, A., Shin, D., Bradley, C. W., Duran-Struuck, R., & Grice, E. A. (2023). Harnessing diversity and antagonism within the pig skin microbiota to identify novel mediators of colonization resistance to methicillin-resistant Staphylococcus aureus. mSphere, 8(4), e00177–23. 10.1128/msphere.00177-23

Wende, M., Osbelt, L., Eisenhard, L., Lesker, T. R., Damaris, B. F., Mutukumarasamy, U., Bielecka, A., d. H. Almási, É., Winter, K. A., Schauer, J., Pfennigwerth, N., Gatermann, S., Schaufler, K., Schlüter, D., Galardini, M., & Strowig, T. (2025). Suppression of gut colonization by multidrug-resistant Escherichia coli clinical isolates through cooperative niche exclusion. Nature Communications, 16, 5426. 10.1038/s41467-025-61327-7

Wertheim, H. F., Melles, D. C., Vos, M. C., van Leeuwen, W., van Belkum, A., Verbrugh, H. A., & Nouwen, J. L. (2005). The role of nasal carriage in Staphylococcus aureus infections. The Lancet Infectious Diseases, 5(12), 751–762. 10.1016/S1473-3099(05)70295-4

Wos-Oxley, M. L., Plumeier, I., von Eiff, C., Taudien, S., Platzer, M., Vilchez-Vargas, R., Becker, K., & Pieper, D. H. (2010). A poke into the diversity and associations within human anterior nare microbial communities. The ISME Journal, 4(7), Article 7. 10.1038/ismej.2010.15

Zipperer, A., Konnerth, M. C., Laux, C., Berscheid, A., Janek, D., Weidenmaier, C., Burian, M., Schilling, N. A., Slavetinsky, C., Marschal, M., Willmann, M., Kalbacher, H., Schittek, B., Brötz-Oesterhelt, H., Grond, S., Peschel, A., & Krismer, B. (2016). Human commensals producing a novel antibiotic impair pathogen colonization. Nature, 535(7613), Article 7613. 10.1038/nature18634

