## Supplementary data for "Inhibition of multidrug-resistant *Staphylococcus aureus* by commensal bacterial species from the human nose"

**Figure S1. Commensal species from the human nose inhibit MRSA strain JE2 in colonisation assays.** Growth of JE2 measured as log(CFU/mL) when co-cultured with combinations of commensal species on BHIA. Each bar represents the mean of four biological replicates ± SE. Statistical comparisons were performed using Dunnett's test. Treatments significantly different from control, $p \leq 0.05$: all treatments except "Ct", "Ne", "Co", "Ca", "Ct,Dp", "Ct,Co", "Ct,Ca", "Dp,Co", "Dp,Ca", "Co,Ca", "Ct,Dp,Ca", "Ct,Co,Ca". Ctrl: no commensal, Sa: *S. aureus*, Sl: *S. lugdunensis*, Ct: *C. tuberculostearicum*, Dp: *D. pigrum*, Ne: *Neisseria sicca*.

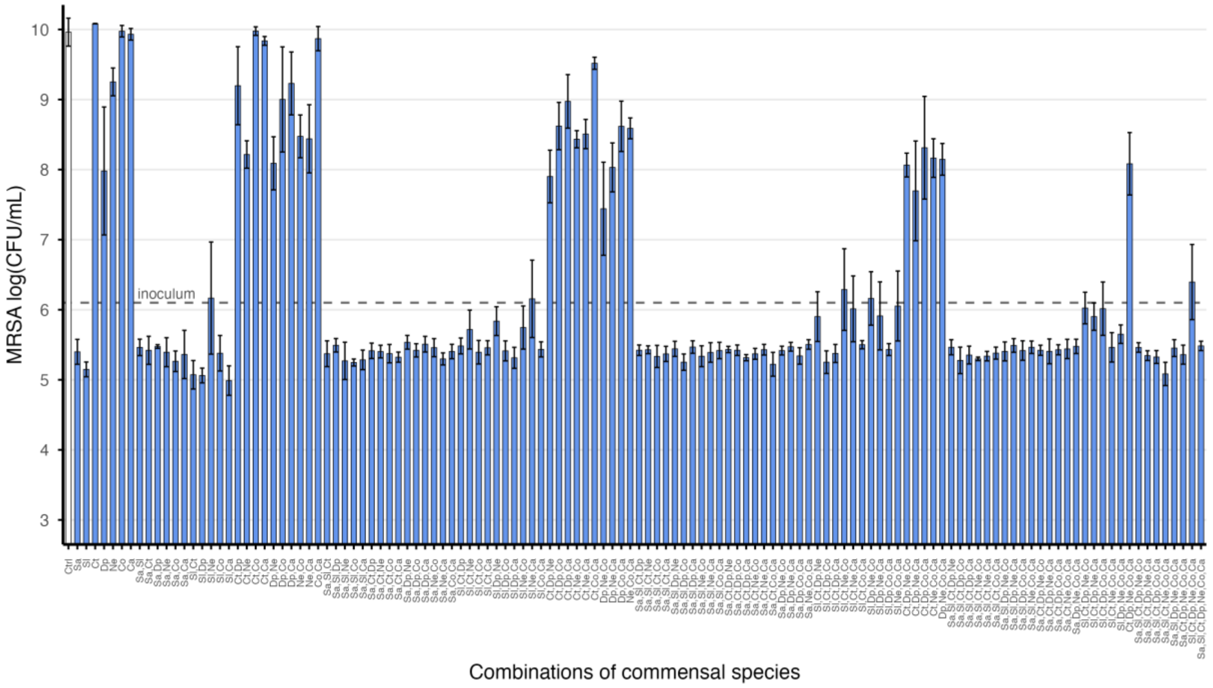

**Figure S2. Effect of Tween 80 on growth of commensals species from the human nose.** Growth of commensal species in presence and absence of 1% Tween 80. Each bar represents the mean of four biological replicates ± SE. Statistical comparisons were performed using unpaired two-tailed t-tests with holm correction for multiple comparisons. Significance: *$padj \leq 0.05$. Sa: *S. aureus*, Sl: *S. lugdunensis*, Ct: *C. tuberculostearicum*, Co: *C. accolens*, Cp: *C. propinquum* Dp: *D. pigrum*, Ne: *Neisseria sicca*, Ca: *C. acnes*.

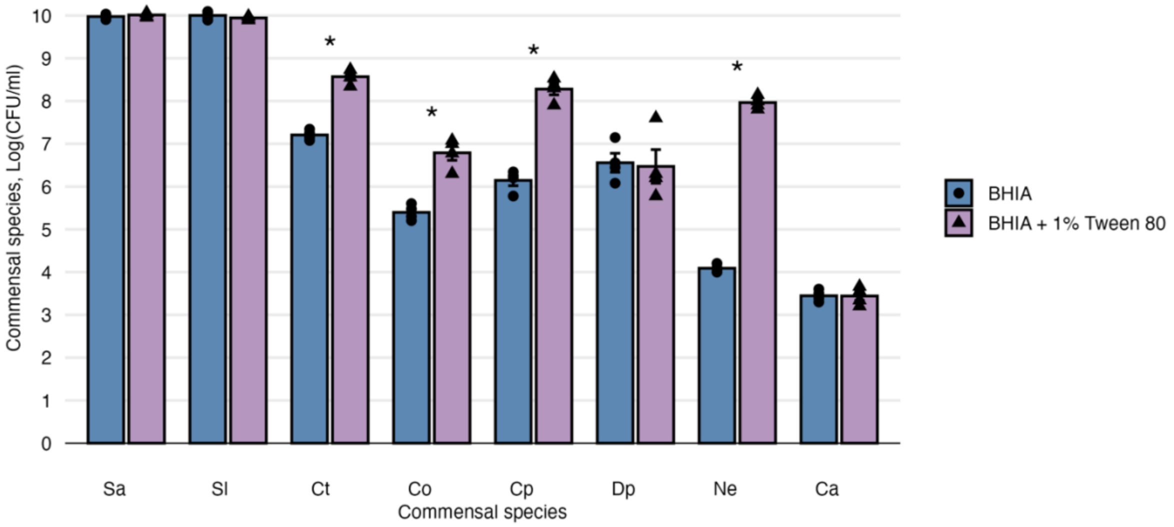

**Figure S3. Doubling time of vancomycin, teicoplanin and daptomycin-adapted strains.** Doubling time of JE2 and vancomycin, teicoplanin and daptomycin-adapted JE2 strains. Cross bars represent mean for six biological replicates. Statistical comparisons were performed using Dunnett's test. Significance: *$p \leq 0.05$. VAN: vancomycin, TEI: teicoplanin, DPC: daptomycin.

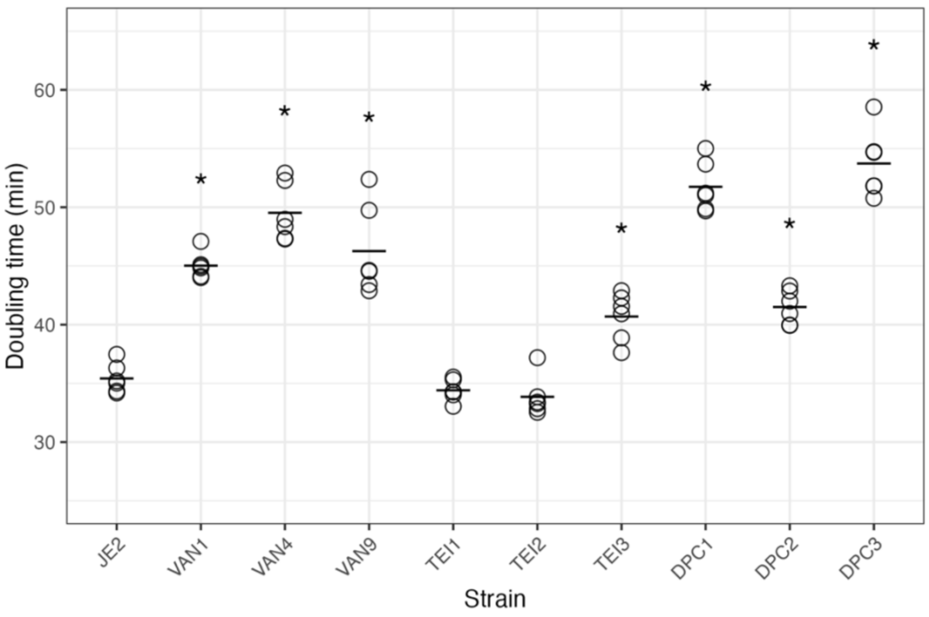

**Figure S4. Doubling time of MRSA strain JE2 adapted to commensal strains.** Doubling time of JE2 wild-type and JE2 passaged in media alone (Ctrl) or with commensal species (Sa: *S. aureus*, Sl: *S. lugdunensis*, Ct: *C. tuberculostearicum*, Dp: *D. pigrum*), with five replicate isolates for each treatment. Cross bars represent the mean for the four replicate assays for each isolate. Statistical comparisons were performed using Dunnett's test. Significance: *$p \leq 0.05$. Ctrl: control, Sa: *S. aureus*, Se: *S. lugdunensis*, Ct: *C. tuberculostearicum*, Dp: *D. pigrum*.

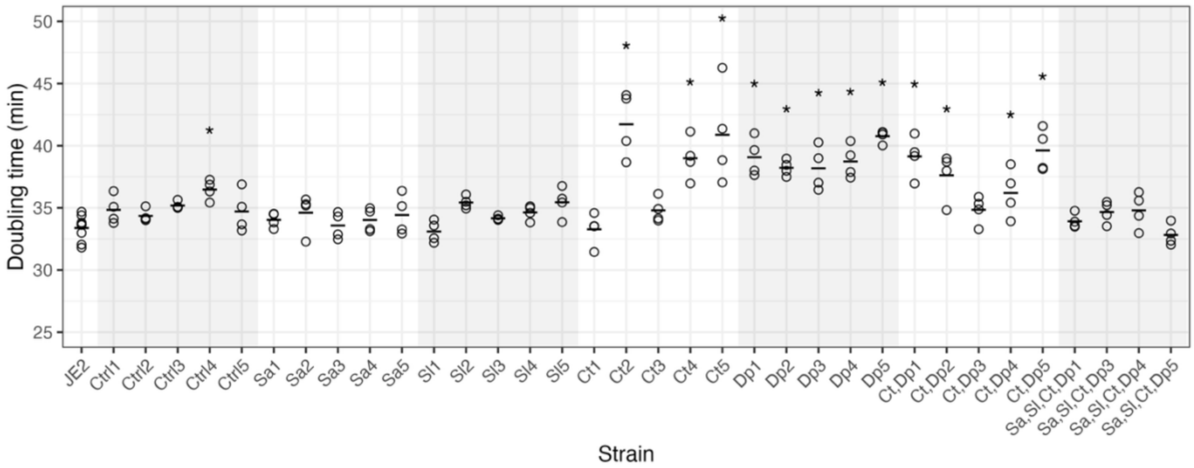

**Figure S5. Inhibition of JE2 transposon mutants during co-culture with commensal species.** Growth of JE2 and transposon mutants alone (Ctrl) and co-cultured with *C. tuberculostearicum* (Ct) alone or together with *D. pigrum* (Dp) on BHIA supplemented with 1% Tween 80. Bars show mean log10(CFU) ± SE; points represent individual replicates. Responses to condition varied among strains (ANOVA, strain × condition interaction, $F_{12, 63} = 5.344$, $p < 0.001$); stars denote significant differences from JE2 within condition (tested by Dunnett's test within each condition, $p < 0.05$).

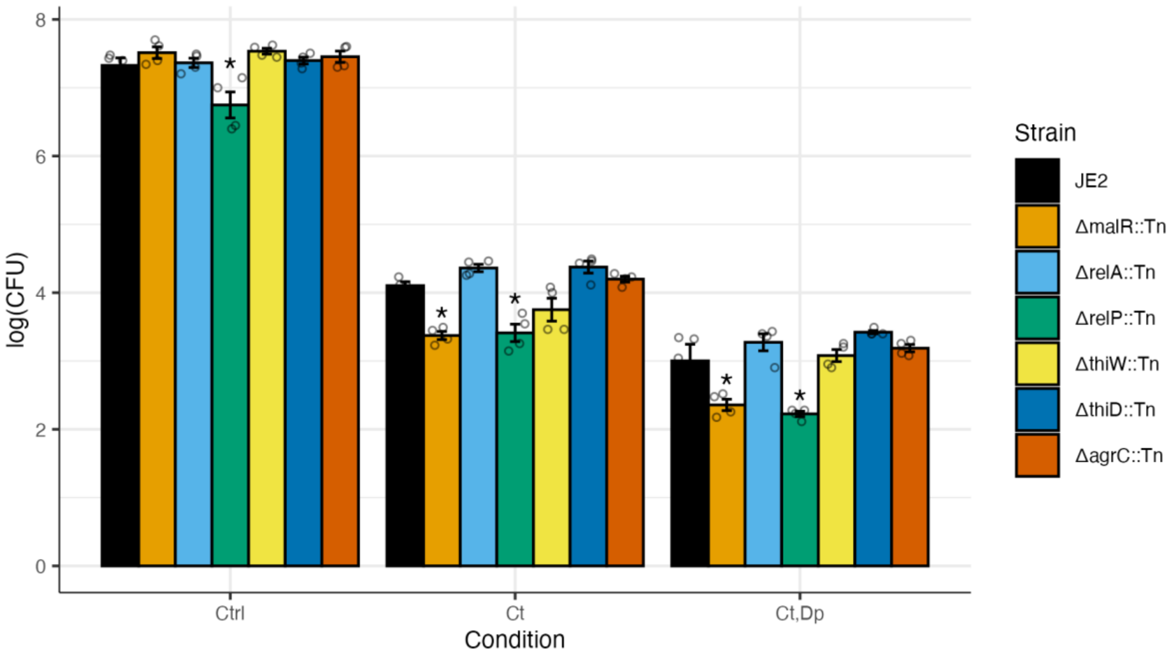

**Table S1. Bacterial strains used in this study.**

| Strain | Species | ID | Description | Source |
|---|---|---|---|---|
| JE2 | *Staphylococcus aureus* | | plasmid-cured derivative of USA300 LAC | Fey *et al.* (2013) |
| VAN1, VAN4, VAN9 | *Staphylococcus aureus* | | laboratory-adapted VISA derivatives of JE2 | Fait *et al.* (2022) (L1, L4, L9) |
| TEI1, TEI2, TEI3 | *Staphylococcus aureus* | | laboratory-adapted derivative of JE2 | This study |
| DPC1, DPC2, DPC3 | *Staphylococcus aureus* | | laboratory-adapted derivative of JE2 | This study |
| JKD6000 (Pair 1) | *Staphylococcus aureus* | | clinical VSSA isolate | Howden *et al.* (2006) |
| JKD6001 (Pair 1) | *Staphylococcus aureus* | | clinical VISA isolate | |
| JKD6004 (Pair 2) | *Staphylococcus aureus* | | clinical VSSA isolate | |
| JKD6005 (Pair 2) | *Staphylococcus aureus* | | clinical VISA isolate | |
| JKD6009 (Pair 3) | *Staphylococcus aureus* | | clinical VSSA isolate | |
| JKD6008 (Pair 3) | *Staphylococcus aureus* | | clinical VISA isolate | |
| JKD6052 (Pair 4) | *Staphylococcus aureus* | | clinical VSSA isolate | |
| JKD6051 (Pair 4) | *Staphylococcus aureus* | | clinical VISA isolate | |
| CI2025 | *Staphylococcus aureus* | Sa | nasal swab | Silvio D. Brugger, University Hospital Zürich |
| CI5657 | *Staphylococcus lugdunensis* | Sl | nasal swab | |
| C62 | *Dolosigranulum pigrum* | Dp | nasal swab | |
| CI5587 | *Corynebacterium tuberculostearicum* | Ct | nasal swab | |
| CI4175 | *Corynebacterium propinquum* | Cp | nasal swab | |
| CI5456 | *Corynebacterium accolens* | Co | nasal swab | |
| CI7291 | *Neisseria sicca* | Ns | nasal swab | |
| CI4246 | *Cutibacterium acnes* | Ca | foreign body | |
| Dp51 | *Dolosigranulum pigrum* | Dp51 | nasal swab | Brülisauer *et al.* (2026) |
| Dp53 | *Dolosigranulum pigrum* | Dp53 | nasal swab | Brülisauer *et al.* (2026) |

**Table S2. Effect of commensal species and interactions on inhibition of MRSA strain JE2.** ANOVA results of the reduced linear model assessing the effects of five commensal species and their interactions on JE2 inhibition measured as log10(CFU/mL). Model reduction was done in a stepwise fashion removing higher order terms using Bayesian Information Criterion (BIC) to compare models (see Methods). This model does not account for differences in species abundance in the different combinations (total inoculum constant across combinations; individual species abundance decreases as species number increases). Asterisks denote significant F-Test ($p \leq 0.05$). Ct: *C. tuberculostearicum*; Dp: *D. pigrum*; Ne: *N. sicca*; Co: *C. accolens*; Ca: *C. acnes*.

| **Source** | **Df** | **SS** | **MS** | **F** | **p** | | **% Var** |
|---|---|---|---|---|---|---|---|
| Ct | 1 | 153.688 | 153.688 | 268.018 | <0.001 | * | 41.469 |
| Dp | 1 | 26.765 | 26.765 | 46.676 | <0.001 | * | 7.222 |
| Ne | 1 | 1.995 | 1.995 | 3.48 | 0.065 | | 0.538 |
| Co | 1 | 60.43 | 60.43 | 105.385 | <0.001 | * | 16.306 |
| Ca | 1 | 0.201 | 0.201 | 0.35 | 0.555 | | 0.054 |
| Ct:Dp | 1 | 0.13 | 0.13 | 0.227 | 0.635 | | 0.035 |
| Ct:Ne | 1 | 4.503 | 4.503 | 7.853 | 0.006 | * | 1.215 |
| Dp:Ne | 1 | 1.565 | 1.565 | 2.729 | 0.101 | | 0.422 |
| Ct:Co | 1 | 37.571 | 37.571 | 65.52 | <0.001 | * | 10.138 |
| Dp:Co | 1 | 0.524 | 0.524 | 0.913 | 0.341 | | 0.141 |
| Ne:Co | 1 | 6.534 | 6.534 | 11.395 | 0.001 | * | 1.763 |
| Dp:Ca | 1 | 0.008 | 0.008 | 0.014 | 0.906 | | 0.002 |
| Ne:Ca | 1 | 0.262 | 0.262 | 0.458 | 0.5 | | 0.071 |
| Ct:Dp:Co | 1 | 4.125 | 4.125 | 7.194 | 0.008 | * | 1.113 |
| Ct:Ne:Co | 1 | 6.009 | 6.009 | 10.48 | 0.002 | * | 1.622 |
| Dp:Ne:Ca | 1 | 2.647 | 2.647 | 4.616 | 0.034 | * | 0.714 |
| Residuals | 111 | 63.65 | 0.573 | | | | 17.174 |

**Table S3. Growth of commensal strains in supernatants of other species.** Growth is expressed as relative $OD_{600nm}$ at 24 h, calculated by dividing growth in strain-derived supernatant by growth in control conditions (supernatant from uninoculated wells). Each column corresponds to the strain used to generate the supernatant (three independent supernatants). Each row corresponds to the strain grown in the indicated supernatant (three biological replicates rep1–rep3). Ctrl: no commensal, Sa: *S. aureus*, Se: *S. lugdunensis*, Ct: *C. tuberculostearicum*, Dp: *D. pigrum* (strains Dp, Dp53 and Dp51), Ns: *N. sicca*. Co: *C. accolens*, Cp: *C. propinquum*, Ca: *Cutibacterium acnes*.

| | | Source of supernatant | | | | | | | | | | |
|---|---|---|---|---|---|---|---|---|---|---|---|---|
| | | Ctrl | Sa | Sl | Ct | Co | Cp | Ns | Ca | Dp | Dp51 | Dp53 |
| Strain growth ($OD_{600nm}$) | Ctrl_rep1 | 0.089 | 0.082 | 0.090 | 0.089 | 0.090 | 0.089 | 0.089 | 0.087 | 0.089 | 0.089 | 0.090 |
| | Ctrl_rep2 | 0.089 | 0.085 | 0.090 | 0.089 | 0.087 | 0.089 | 0.089 | 0.088 | 0.089 | 0.090 | 0.086 |
| | Ctrl_rep3 | 0.088 | 0.090 | 0.088 | 0.089 | 0.089 | 0.089 | 0.089 | 0.088 | 0.090 | 0.089 | 0.086 |
| | Sa_rep1 | 0.181 | 0.089 | 0.104 | 0.123 | 0.135 | 0.119 | 0.138 | 0.174 | 0.225 | NA | NA |
| | Sa_rep2 | 0.193 | 0.093 | 0.102 | 0.116 | 0.129 | 0.114 | 0.130 | 0.276 | 0.241 | NA | NA |
| | Sa_rep3 | 0.208 | 0.089 | 0.101 | 0.115 | 0.122 | 0.113 | 0.139 | 0.272 | 0.246 | NA | NA |
| | Sl_rep1 | 0.183 | 0.094 | 0.092 | 0.103 | 0.108 | 0.105 | 0.114 | 0.174 | 0.179 | NA | NA |
| | Sl_rep2 | 0.184 | 0.094 | 0.092 | 0.113 | 0.107 | 0.102 | 0.112 | 0.179 | 0.196 | NA | NA |
| | Sl_rep3 | 0.189 | 0.092 | 0.093 | 0.106 | 0.099 | 0.104 | 0.113 | 0.160 | 0.215 | NA | NA |
| | Ct_rep1 | 0.147 | 0.090 | 0.090 | 0.089 | 0.092 | 0.093 | 0.095 | 0.145 | 0.174 | 0.207 | 0.251 |
| | Ct_rep2 | 0.166 | 0.092 | 0.091 | 0.092 | 0.090 | 0.093 | 0.091 | 0.302 | 0.202 | 0.218 | 0.254 |
| | Ct_rep3 | 0.179 | 0.091 | 0.094 | 0.094 | 0.090 | 0.094 | 0.093 | 0.204 | 0.219 | 0.209 | 0.207 |
| | Co_rep1 | 0.140 | 0.090 | 0.090 | 0.096 | 0.090 | 0.095 | 0.113 | 0.122 | 0.187 | 0.230 | 0.249 |
| | Co_rep2 | 0.166 | 0.092 | 0.091 | 0.104 | 0.091 | 0.093 | 0.108 | 0.278 | 0.142 | 0.213 | 0.229 |
| | Co_rep3 | 0.160 | 0.088 | 0.092 | 0.104 | 0.089 | 0.091 | 0.108 | 0.267 | 0.291 | 0.226 | 0.237 |
| | Cp_rep1 | 0.242 | 0.134 | 0.138 | 0.173 | 0.164 | 0.099 | 0.139 | 0.205 | 0.241 | 0.234 | 0.235 |
| | Cp_rep2 | 0.252 | 0.119 | 0.136 | 0.197 | 0.138 | 0.105 | 0.139 | 0.224 | 0.251 | 0.245 | 0.246 |
| | Cp_rep3 | 0.234 | 0.114 | 0.135 | 0.120 | 0.123 | 0.102 | 0.147 | 0.177 | 0.274 | 0.186 | 0.174 |
| | Ns_rep1 | 0.103 | 0.092 | 0.093 | 0.088 | 0.091 | 0.094 | 0.092 | 0.123 | 0.089 | NA | NA |
| | Ns_rep2 | 0.162 | 0.103 | 0.094 | 0.089 | 0.091 | 0.092 | 0.090 | 0.100 | 0.123 | NA | NA |
| | Ns_rep3 | 0.140 | 0.086 | 0.110 | 0.093 | 0.090 | 0.092 | 0.090 | 0.110 | 0.173 | NA | NA |
| | Dp_rep1 | 0.089 | 0.082 | 0.086 | 0.091 | 0.088 | 0.093 | 0.091 | 0.087 | 0.087 | NA | NA |
| | Dp_rep2 | 0.090 | 0.088 | 0.089 | 0.096 | 0.088 | 0.091 | 0.089 | 0.089 | 0.087 | NA | NA |
| | Dp_rep3 | 0.088 | 0.089 | 0.089 | 0.092 | 0.090 | 0.094 | 0.089 | 0.090 | 0.085 | NA | NA |
| | Ca_rep1 | 0.097 | 0.085 | 0.089 | 0.092 | 0.090 | 0.093 | 0.092 | 0.090 | 0.103 | NA | NA |
| | Ca_rep2 | 0.094 | 0.093 | 0.092 | 0.088 | 0.090 | 0.092 | 0.089 | 0.089 | 0.093 | NA | NA |
| | Ca_rep3 | 0.092 | 0.094 | 0.092 | 0.094 | 0.090 | 0.095 | 0.091 | 0.090 | 0.094 | NA | NA |

**Table S4. Susceptibility of daptomycin and teicoplanin-adapted strains.** MIC were measured using antibiotic strips. MIC Breakpoints for Daptomycin and Teicoplanin and vancomycin are >1 and >2, respectively (EUCAST, 2024).

| | MIC (mg/L) | |
|---|---|---|
| | Daptomycin | Teicoplanin |
| JE2 | 0.5 | 2 |
| DPC1 | 4 | 3.5 |
| DPC2 | 4 | 4.5 |
| DPC3 | 4 | 4.5 |
| TEI1 | 0.5 | 4.5 |
| TEI2 | 1.5 | 5 |
| TEI3 | 1 | 5 |

**Table S5. Mutations acquired during adaptation of MRSA strain JE2 to teicoplanin and daptomycin.** The MRSA strain JE2 was grown on increasing concentrations of teicoplanin and daptomycin in ten independent parallel populations; a single colony isolate from three of the ten populations for each antibiotic was then selected for this study (labelled at left).

| | TYPE | LOCUS_TAG | GENE | EFFECT | PRODUCT |
|---|---|---|---|---|---|
| TEI1 | snp | SAUSA300_0021 | *walK* | Val15Ala | sensory box histidine kinase |
| TEI2 | snp | SAUSA300_1867 | *vraT* | Pro174Gln | conserved hypothetical protein |
| TEI3 | snp | SAUSA300_0527 | *rpoB* | Gly224Asp | DNA-directed RNA polymerase, beta subunit |
| | snp | SAUSA300_0995 | | Gly11Trp | dihydrolipoamide acetyltransferase |
| DPC1 | snp | intergenic | | | |
| | snp | SAUSA300_1255 | *mprF* | Asn352Lys | oxacillin resistance-related MprF protein |
| | snp | SAUSA300_2071 | *prmC* | Gly89Arg | modification methylase, HemK family |
| DPC2 | snp | intergenic | | | |
| | del | SAUSA300_1867 | *vraT* | Tyr24fs | conserved hypothetical protein |
| DPC3 | snp | SAUSA300_0214 | | Glu52Glu | conserved hypothetical protein |
| | snp | SAUSA300_1176 | *pgsA* | Phe76Ile | CDP-diacylglycerol--glycerol-3-phosphate 3-phosphatidyltransferase |
| | snp | SAUSA300_1295 | | Ser31Ala | cold shock protein, CSD family |

**Table S6. Antibiotic susceptibility of commensal-adapted JE2 isolates.** MIC of commensal-adapted JE2 isolates were measured using antibiotic strips. MIC Breakpoints for Daptomycin and Teicoplanin and vancomycin are >1 and >2, respectively (EUCAST, 2024). JE2 adapted to commensal treatments: Ctrl: control (no commensal, media adaptation), Sa: *S. aureus*, Se: *S. lugdunensis*, Ct: *C. tuberculostearicum*, Dp: *D. pigrum*, Ct,Dp and Sa,Sl,Ct,Dp.

| | MIC (mg/L) | |
|---|---|---|
| | Daptomycin | Teicoplanin |
| JE2 | 1 | 1.5 |
| Ctrl1 | 1 | 1 |
| Ctrl2 | 1 | 1 |
| Ctrl3 | 1 | 1.5 |
| Ctrl4 | 1 | 1.5 |
| Ctrl5 | 1 | 1.5 |
| Sa1 | 1 | 1.5 |
| Sa2 | 0.75 | 1.5 |
| Sa3 | 1 | 2 |
| Sa4 | 0.75 | 1.5 |
| Sa5 | 1 | 1.5 |
| Sl1 | 1 | 1.5 |
| Sl2 | 0.75 | 1 |
| Sl3 | 0.75 | 1.5 |
| Sl4 | 0.5 | 1 |
| Sl5 | 1 | 1.5 |
| Ct1 | 1 | 1.5 |
| Ct2 | 0.75 | 0.75 |
| Ct3 | 0.75 | 1 |
| Ct4 | 0.75 | 1 |
| Ct5 | 1 | 1 |
| Dp1 | 1 | 1.5 |
| Dp2 | 0.75 | 1 |
| Dp3 | 1 | 1.5 |
| Dp4 | 1 | 1.5 |
| Dp5 | 1 | 1.5 |
| Ct,Dp1 | 1 | 1 |
| Ct,Dp2 | 1 | 1 |
| Ct,Dp3 | 1 | 1 |
| Ct,Dp4 | 0.75 | 1 |
| Ct,Dp5 | 1 | 1 |
| Sa,Sl,Ct,Dp1 | 1 | 1.5 |
| Sa,Sl,Ct,Dp2 | 0.75 | 1.5 |
| Sa,Sl,Ct,Dp3 | 1 | 1.5 |
| Sa,Sl,Ct,Dp4 | 1 | 1.5 |

**Table S7. Genetic changes in JE2 isolates following adaptation to commensals.**

| | TYPE | LOCUS_TAG | GENE | EFFECT | PRODUCT |
|---|---|---|---|---|---|
| **Ctrl1** | del | SAUSA300_1991 | *agrC* | Ile52del | accessory gene regulator protein C |
| **Ctrl2** | - | | | | |
| **Ctrl3** | snp | SAUSA300_1591 | *apt* | Tyr162Asp | adenine phosphoribosyltransferase |
| **Ctrl4** | snp | SAUSA300_1591 | *apt* | Tyr162Asp | adenine phosphoribosyltransferase |
| **Ctrl5** | - | | | | |
| **Ct1** | ins | SAUSA300_1457 | *malR* | Leu83fs | maltose operon transcriptional repressor |
| **Ct2** | snp | SAUSA300_1590 | *relA* | Thr320Ile | GTP pyrophosphokinase |
| **Ct3** | snp | SAUSA300_0978 | *thiW* | Glu295* | ABC transporter, ATP-binding protein |
| | snp | SAUSA300_1590 | *relA* | Gly81Ser | GTP pyrophosphokinase |
| **Ct4** | del | SAUSA300_1590 | *relA* | Gln679_Ser683delinsPro | GTP pyrophosphokinase |
| **Ct5** | snp | SAUSA300_0562 | *thiD* | Gly248Asp | phosphomethylpyrimidine kinase |
| | snp | SAUSA300_1590 | *relA* | Asp141Tyr | GTP pyrophosphokinase |
| **CtDp1** | snp | SAUSA300_2446 | *relP* | Ser21Ile | conserved hypothetical protein |
| **CtDp2** | snp | SAUSA300_1590 | *relA* | Glu676* | GTP pyrophosphokinase |
| **CtDp3** | snp | SAUSA300_2446 | *relP* | His69Gln | conserved hypothetical protein |
| | snp | | | 2841979G>T | |
| **CtDp4** | snp | SAUSA300_0562 | *thiD* | Trp249* | phosphomethylpyrimidine kinase |
| | snp | SAUSA300_1590 | *relA* | Ala80Val | GTP pyrophosphokinase |
| **CtDp5** | snp | SAUSA300_1590 | *relA* | Arg132Ser | GTP pyrophosphokinase |
